# A deep generative decoder predicts microRNA expression from bulk and single-cell mRNA profiles

**DOI:** 10.64898/2026.05.29.727918

**Authors:** Farhad Zamani, Asta Mannstaedt Rasmussen, Viktoria Schuster, Mathilde Hartvig Diekema, Anders Krogh, Jakob Skou Pedersen

## Abstract

MicroRNAs (miRNAs) are key post-transcriptional regulators, yet standard bulk and single-cell RNA-seq do not capture them, leaving this regulatory layer invisible in most transcriptomic data. We present miDGD, a deep generative decoder that jointly models paired mRNA and miRNA profiles through a shared latent representation, enabling miRNA expression to be predicted from mRNA alone. Trained on tumors (TCGA), healthy tissues (GTEx), and human cell lines, miDGD recovers hundreds of miRNAs in held-out tumors (mean Spearman ρ = 0.56), captures both tissue-specific and ubiquitous miRNAs, and preserves known miRNA–target repression and host-gene co-expression. Without label supervision, its latent space separates 32 cancer types (80% accuracy). Predictions remain stable at single-cell-like sparsity and transfer across datasets—from tumors to healthy tissues and from bulk to single cells—where miDGD outperforms existing supervised and activity-inference methods. miDGD thus unlocks miRNA regulation in the vast body of existing mRNA-only data, including single-cell datasets.

**GRAPHICAL ABSTRACT:** 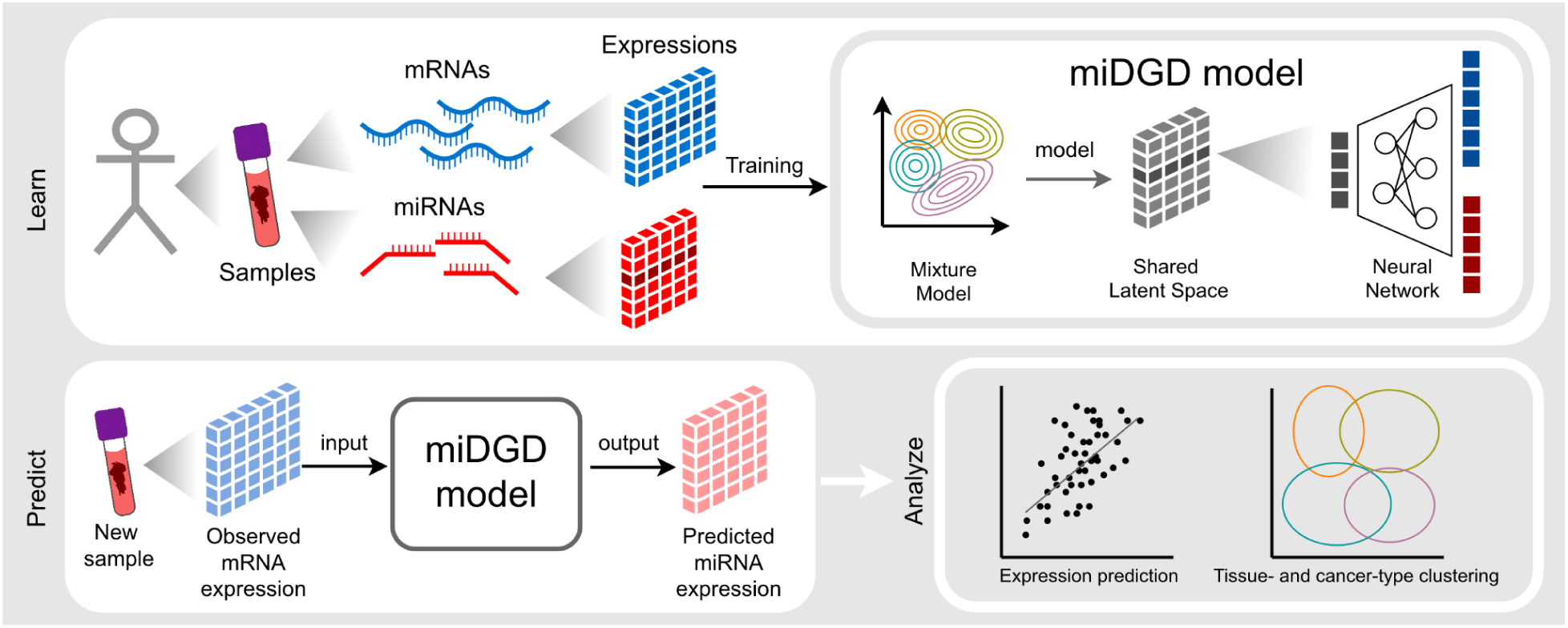

## INTRODUCTION

MicroRNAs (miRNAs) are small non-coding RNAs that regulate gene expression post-transcriptionally by promoting target mRNA degradation, thereby shaping cellular transcriptional programs^1–3^. Mature miRNAs, either the 5p or 3p arm, guide the RNA-induced silencing complex (RISC), facilitating mRNA translation repression through target recognition followed by destabilization and degradation (Fig. 1a)^1,3^. Bulk tissue studies have shown that miRNA regulation is highly tissue-specific^4,5^, contributing to tissue identity and homeostasis, and is perturbed in multiple diseases^6,7^, including during cancer development^7–9^. Here, miRNAs exhibit both oncogenic and tumor-suppressor capabilities^7^, making them candidate biomarkers and therapeutic targets; for example, miR-9 is deregulated in glioma^10,11^, miR-21 is widely upregulated during cancer progression^12,13^, and miR-122 is down-regulated in hepatocellular carcinoma^14,15^. While miRNAs are expected to exhibit highly spatio-temporal expression patterns and perform cell-type-specific gene regulation, they remain poorly studied at the single-cell level, with only recent protocols enabling low-throughput and non-routine detection^16–19^.

To mitigate current limitations in single-cell miRNA detection, several computational methods have been developed for indirect inference^20–24^. Unsupervised approaches infer miRNA activity from the depletion of target genes relative to all other expressed genes, with latest methods including miTEA-HiRes^23^ and bayesReact^24^. In contrast, supervised models can learn complex relationships among miRNAs, their target genes, transcription factors, and host gene expression to directly predict miRNA expression. miRSCAPE was the first to implement this strategy, using paired bulk mRNA–miRNA data and extreme gradient boosting (XGBoost)^25,26^ to infer miRNA levels from single-cell mRNA expression. However, because it fits a separate model for each miRNA, it cannot leverage co-regulation or shared structure across miRNAs, which may limit both training sample efficiency and generalization across datasets. In addition, miRNA prediction from single-cell RNA-seq (scRNA-seq) remains challenging due to distributional and technical differences from bulk RNA-seq, including library preparation, isolated cells rather than cell mixtures, sparse expression profiles, dropout rates, and limited benchmarking data for evaluation^19,27^. While atlas-scale models^28–30^ have advanced single-cell analysis more broadly, they are not designed for miRNA prediction from mRNA, where available training data are orders of magnitude smaller.

Deep Generative Decoders (DGDs) are well-suited to this setting^31^. By learning sample-specific latent variables directly as trainable parameters without an encoder, they require fewer parameters and are substantially more data-efficient than encoder–decoder architectures. A mixture-based prior further yields a multimodal latent space that captures discrete subpopulations and continuous variation across tissues and cancer types. For paired mRNA–miRNA data, this enables a shared low-dimensional representation that captures joint structure across both modalities even in comparatively small training cohorts. The DGD framework has previously been applied to bulk and single-cell transcriptomics and to joint transcriptomic and chromatin-accessibility modeling, demonstrating biologically interpretable latent structure and robust cross-dataset generalization^31–33^.

In this work, we introduce miDGD, a Deep Generative Decoder that learns a shared low-dimensional representation of paired mRNA-miRNA expression profiles and uses it to predict miRNA expression from gene expression alone. Using The Cancer Genome Atlas (TCGA)^34^ as a core pan-cancer training set together with the Genotype-Tissue Expression (GTEx) project^35^, R2 RNA Atlas^36^, and single-cell human cell line data^18,19^, we demonstrate that miDGD reconstructs biologically meaningful expression patterns for hundreds of miRNAs in multiple independent cohorts, preserving miRNA–target repression and host-gene relationships. Uninformed of cancer-type labels, the learned latent space recovers cancer-type identity across the TCGA pan-cancer cohort and allows label prediction for new samples. We further show that miDGD generalizes to datasets not seen during training and maintains robust performance under strong downsampling that mimics single-cell sparsity. Finally, we apply miDGD to scRNA-seq from human cell lines to impute miRNA expression at single-cell resolution, where it outperforms existing supervised and unsupervised methods. Together, these findings demonstrate that miRNAs, post-transcriptional regulators, are sufficiently associated with mRNA expression to be reconstructed from conventional transcriptomic data, establishing deep generative modeling as a practical route to analyze miRNA regulation in existing public data.

## RESULTS

### miDGD Overview

Gene regulation is best understood through multimodal measurements, yet many regulatory layers remain unobserved in the same samples or cells. miRNAs are a prominent example; despite their central post-transcriptional roles, they remain largely unmeasured in individual cells due to experimental constraints^16,19^. Joint modelling of mRNA and miRNA expression (Fig. 1b) can enable prediction of miRNA expression in settings where it would otherwise be unobserved. A shared low-dimensional representation can capture structural associations among miRNAs, their target mRNAs, host genes, and other co-regulators such as transcription factors.

**Figure 1:**
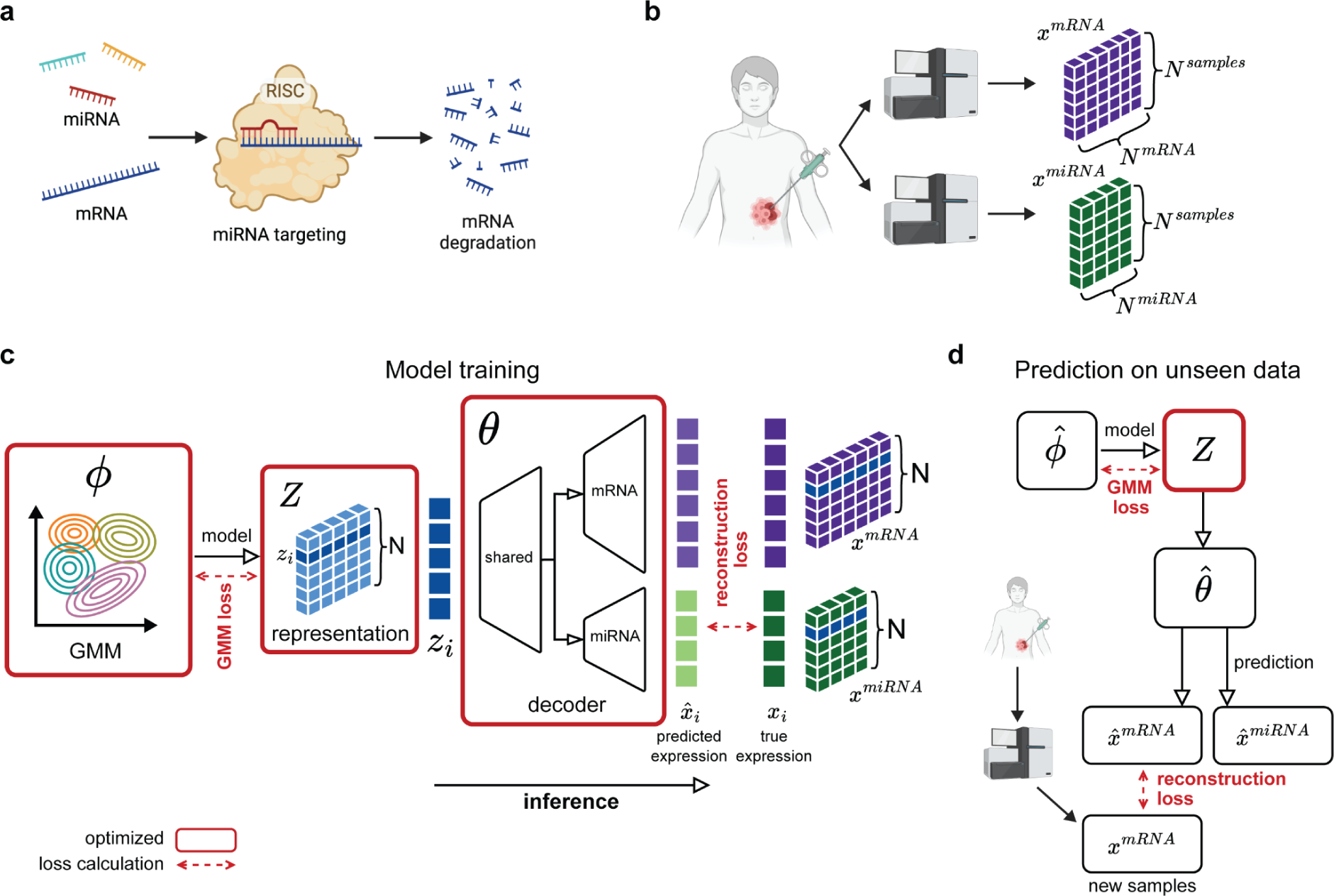
Overview of miDGD and data input. **a**, microRNA (miRNA) binds to target mRNA to regulate expression and repress translation. **b**, Paired mRNA and miRNA expression profiled from the same samples or cells. **c**, Schematic of miDGD model training and data generative process. The model consists of representation *Z*, GMM **φ**, and decoder θ, which are updated during optimization (depicted in red box). GMM, Gaussian mixture model. **d**, Application of the trained miDGD to predict miRNA expression from a new set of mRNA expression data. During this step, only the latent representation *Z* of the test samples is optimized, while the GMM **φ̂** and decoder θ̂ parameters are kept fixed.

To do so, we developed miDGD by adapting the multiDGD framework^32^ to paired mRNA and miRNA count data. miDGD learns a low-dimensional latent representation for each sample together with a shared decoder that maps the representation to both modalities. A Gaussian mixture model (GMM) prior facilitates latent-space structure that captures major expression programs across cancer and tissue types. Because miDGD is encoderless, the model parameters of the decoder, θ, and the prior, **φ**, as well as the sample-specific representations (*z_i_* ∈*Z*) are all optimized directly via maximum a posteriori estimation under count-based negative binomial likelihoods during training (Fig. 1c). We trained miDGD models on matched mRNA–miRNA expression profiles from bulk and single-cell RNA-seq datasets, split into training, validation, and test, with hyperparameters optimized for each model–dataset combination (see Methods).

For new data in which only mRNA is observed, miDGD infers a latent representation in two steps (Fig. 1d). First, each new sample (or cell) is initialized at the component mean whose decoded mRNA profile best matches the observed mRNA counts. Second, with decoder and GMM parameters fixed, only the sample representation is refined by gradient-based optimization to minimize the mRNA reconstruction loss. The optimized representation is then passed through the shared decoder to generate the corresponding miRNA profile (see Methods for details).

### Representations Recover Cancer Type Clusters

To evaluate whether miDGD learns meaningful representations, we examined how samples are organized in latent space. Using the pan-cancer TCGA set (Fig. 2a; Supplementary Table 1), we first visualized the joint representation learned from both mRNA and miRNA training profiles (Fig. 2b). Samples form well-separated clusters by cancer type, and GMM component means lie near the centers of these clusters (Fig. 2b,c), indicating that the shared latent space recovers known cancer type structure in the data. Despite the model being trained unsupervised with respect to cancer type labels, the sample-to-component clustering shows that clusters reflect known cancer types (Adjusted Rand Index, ARI, of 0.39; Fig. 2c).

**Figure 2:**
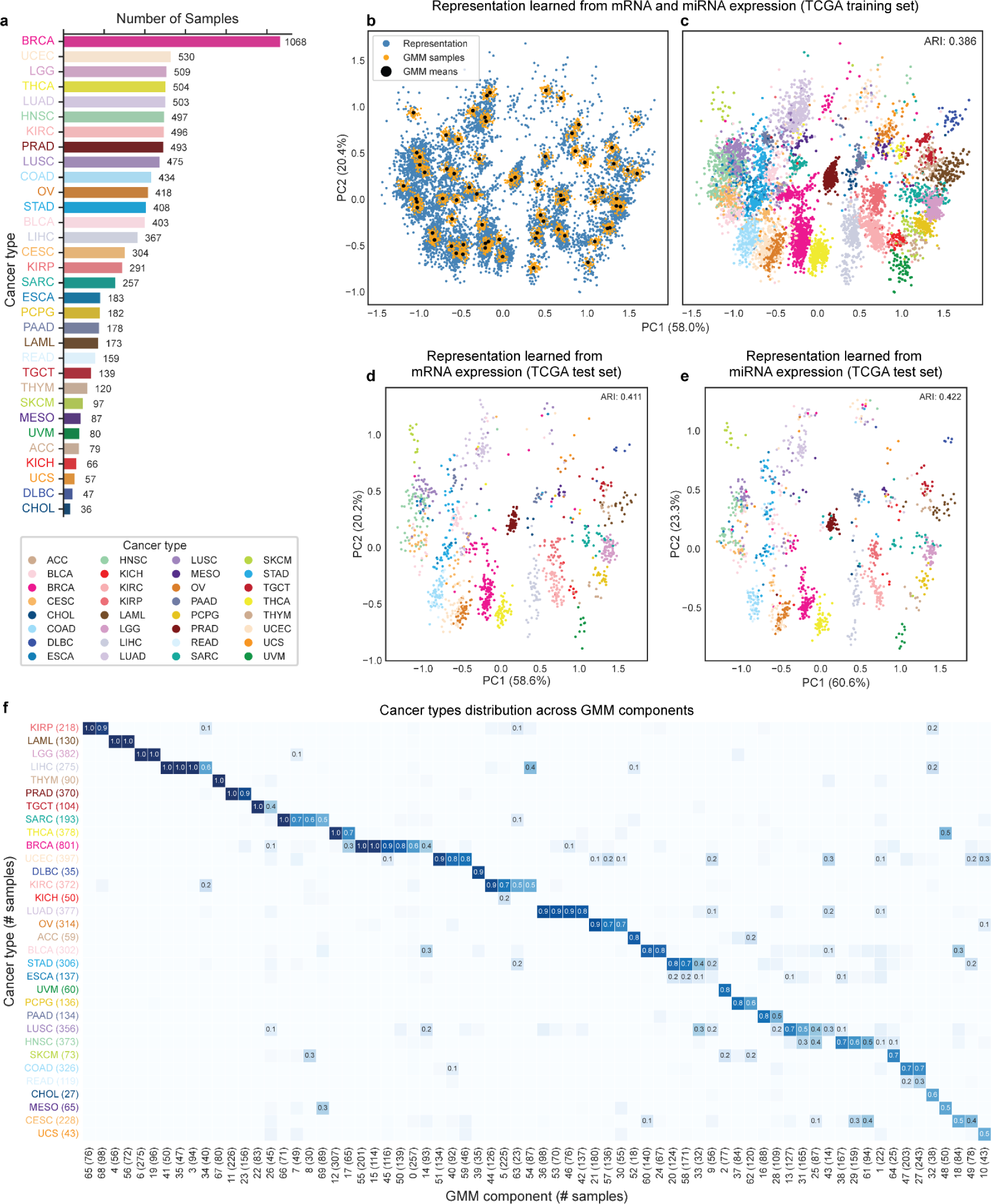
Latent representations capture cancer type structure. **a**, Number of TCGA samples per cancer type. **b**, PCA of latent space for TCGA training samples, showing sample representations (blue) together with GMM samples (orange) and component means (black). **c**, Analogous with panel b, depicting sample representations coloured by cancer type. **d**, Latent representations inferred on the held-out test set using either mRNA alone or, **e**, miRNA alone. **f**, Matrix summarizing the fraction of samples from each cancer type assigned to a given Gaussian component (columns sum to one).

Further evaluation showed that samples inferred from a single modality still embed meaningfully into the learned latent space structured by cancer type. On the held-out test set, we inferred latent representations either from mRNA alone or miRNA alone using the trained decoder and GMM (Fig. 2d,e). In both cases, samples continued to cluster by cancer type, showing that miDGD supports consistent placement of new samples in the latent space even when only one modality is available (Supplementary Fig. 1). Using the fixed GMM components to predict cancer type by majority vote, mRNA-only inference achieves 80% accuracy across 32 cancer types (ARI = 0.41, homogeneity = 0.84, completeness = 0.67). Six cancer types were recovered perfectly and over 80% of types exceed 85% accuracy. Meanwhile, instances with low accuracy were mainly due to assignment of samples to similar cancer types from the same or adjacent tissue of origin (Supplementary Table 2). miRNA-only inference achieves comparable clustering quality (ARI = 0.42) and prediction accuracy (77%).

To assess how the joint representation reflects cancer type, we visualized the distribution of cancer types across GMM components (Fig. 2f). For each component, we then summarized the fraction of its samples belonging to the most frequent cancer type as a purity score. High-purity components indicate cancer-type-specific regions of the latent space. For example, Kidney renal papillary cell carcinoma (KIRP), Acute myeloid leukemia (LAML), and Lower-grade glioma (LGG) are each almost entirely captured by two highly pure components (Fig. 2f).

Some cancer types are dispersed across multiple components, suggesting heterogeneous within-type structure. For instance, Breast invasive carcinoma (BRCA) and Liver hepatocellular carcinoma (LIHC), are represented by multiple components, consistent with within-type molecular substructure classified by PAM50 mRNA subtypes in BRCA^37^ and iCluster subtypes in LIHC^38^ (Supplementary Fig. 2). In contrast, other cancer types show overlapping representation in latent space: Colon adenocarcinoma (COAD) and Rectum adenocarcinoma (READ) samples are mainly represented (>90%) in the same two components, consistent with the proximity of their primary tumor sites and similarities in their transcriptomic profiles, and Kidney Chromophobe (KICH) is consistently assigned with Kidney renal clear cell carcinoma (KIRC), likely reflecting shared kidney tissue-of-origin profiles^39,40^. Overall, miDGD generates a well-organized latent space that captures both cancer-type identity and within-cancer-type variation.

### miDGD Recovers Both Cancer-Type-Specific and Ubiquitous microRNA Expression Profiles

To evaluate miDGD’s ability to recover biologically meaningful miRNA patterns from mRNA data, we first assessed its ability to predict miRNA expression profiles from mRNA counts in TCGA primary tumor test samples (n = 1,205). A positive correlation is obtained between observed and predicted miRNA expressions (mean Pearson *r* = 0. 57, mean Spearman ρ = 0. 56; Fig. 3a,b, Supplementary Fig. 23a,b). Considering highly correlated miRNAs, we find that miDGD can reconstruct both cancer-type-specific and ubiquitous miRNA profiles, with examples including the neuronal miR-9-5p and oncogenic miR-21-5p, respectively (Fig. 3c,d). Thus, the trained model captures useful relationships between mRNAs and individual miRNAs across samples from various cancer types. Some miRNAs remained difficult to predict, potentially reflecting a weak relationship between their mature transcript levels and bulk mRNA expression, due to additional post-transcriptional and cell-type-specific regulatory layers not directly captured by mRNA abundance alone. Many poorly correlated miRNAs also exhibit low expression levels, which likely contributes to weaker correlation (Supplementary Fig. 23c,d). However, a majority of miRNAs show a moderate to high correlation between observed and predicted expression (88% have *r* > 0. 3, median *r* = 0. 60).

**Fig. 3:**
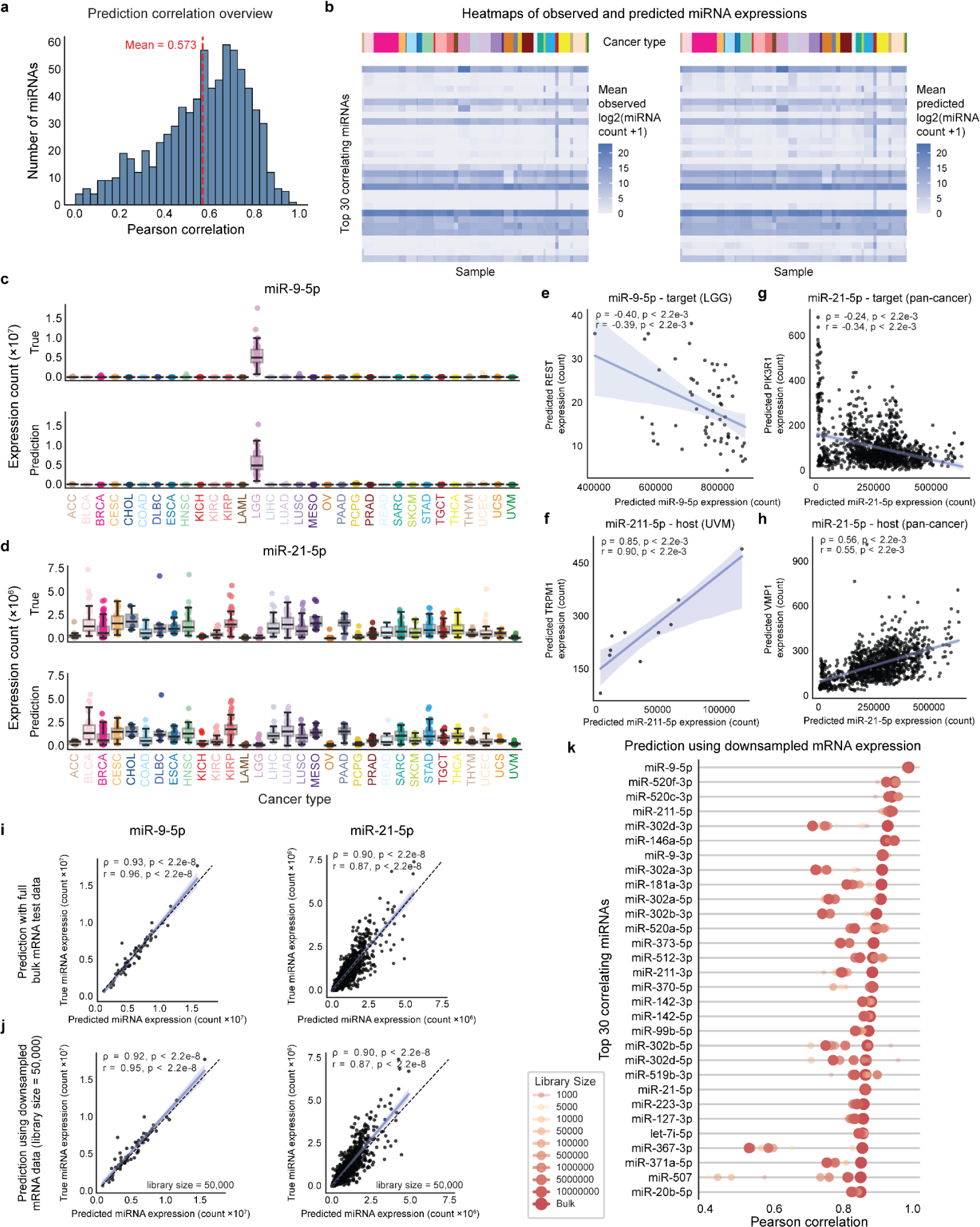
miDGD accurately recovers miRNA expression profiles in TCGA primary tumors. **a**, Pearson correlation between observed and predicted miRNA expression (n = 755) across the TCGA test set samples (n = 1,205). **b**, Heatmaps depicting mean miRNA expression (left) and predictions (right) within each cancer type. Cancer type ordering matches panel c, and miRNAs match panel k. **c**, True and predicted expression of miR-9-5p and, **d**, miR-21-5p across cancer types. **e**, Association between predicted expression of miR-9-5p and predicted expression of known target gene REST in LGG; **f**, between predicted miR-211-5p and its melanocyte-specific host gene TRPM1 in UVM; **g**, between predicted miR-21-5p and its target gene PIK3R1; **h**, between predicted miR-21-5p and its host gene VMP1. **i**, True versus predicted expression of miR-9-5p and miR-21-5p, based on full-depth mRNA expression data, and **j**, downsampled mRNA expression data (library size of 50,000). **k**, Pearson correlations of the top 30 best-predicted miRNAs in the TCGA test data and corresponding semi-synthetic datasets with downsampled gene expression to fixed library sizes.

To examine individual miRNAs in more detail, we focused on miR-9-5p and miR-21-5p, which are tissue-enriched and broadly expressed, respectively (Fig. 3c,d). miR-9-5p is accurately recovered in brain tumors, where it is highly expressed, and remains near zero elsewhere, whereas miR-21-5p is well predicted across many tumor types (miR-9-5p *r* = 0. 93, miR-21-5p *r* = 0. 90; Fig. 3i).

Furthermore, we considered whether miDGD’s predictions preserve known miRNA–mRNA relationships. We correlated predicted miRNA abundances with predicted mRNA expressions of well-known target and host genes across TCGA samples. Predicted miR-9-5p levels were negatively associated with its target gene REST^41^(Fig. 3e), and predicted miR-21-5p levels were negatively associated with the target gene PIK3R1^42^ (Fig. 3f), consistent with the expected repressive effect of these miRNAs on their targets. In addition, miDGD reproduced positive associations with host genes, including the melanocyte-specific TRPM1 for miR-211-5p in uveal melanoma (UVM)^43^ and miR-21-5p with host gene VMP1^44,45^(Fig. 3g,h). Together, these examples show that miDGD not only recovers global miRNA expression patterns but also maintains biologically meaningful miRNA–target and host–miRNA relationships.

Since miDGD is intended for a broad range of coverage levels, including sparse single-cell data, we assessed its robustness to input sparsity by downsampling mRNA counts to generate increasingly sparse datasets, with some depths mimicking scRNA-seq (< 200,000 counts per sample). These were used as input to predict miRNA expression. Correlations between observed and predicted miRNA expression remained high across increasing sparsity (Fig. 3j). Considering the 30 best-predicted miRNAs, performance was stable over several orders of magnitude of downsampling, down to 1,000 counts per sample (Fig. 3k; Supplementary Fig. 23e), indicating that the learned mRNA-to-miRNA mapping is robust to sparse input and that strongly predictable miRNAs can still be reliably inferred at low depth.

### Cross-Dataset Performance on Healthy Tissue Samples

Generalization to new, independent datasets remains challenging for specialized models due to differences in biological context and technical variation, potentially leading to shifted or out-of-distribution observations^46^. To evaluate both within- and cross-dataset miRNA prediction, we trained miDGD on different training sets, including healthy tissue samples from GTEx (n = 10,778)^35,47^, which recently added small RNA-seq to its total RNA-seq resource (Fig. 4a), and assessed the prediction of 728 miRNAs in a held-out test set (n = 2,310).

**Fig. 4:**
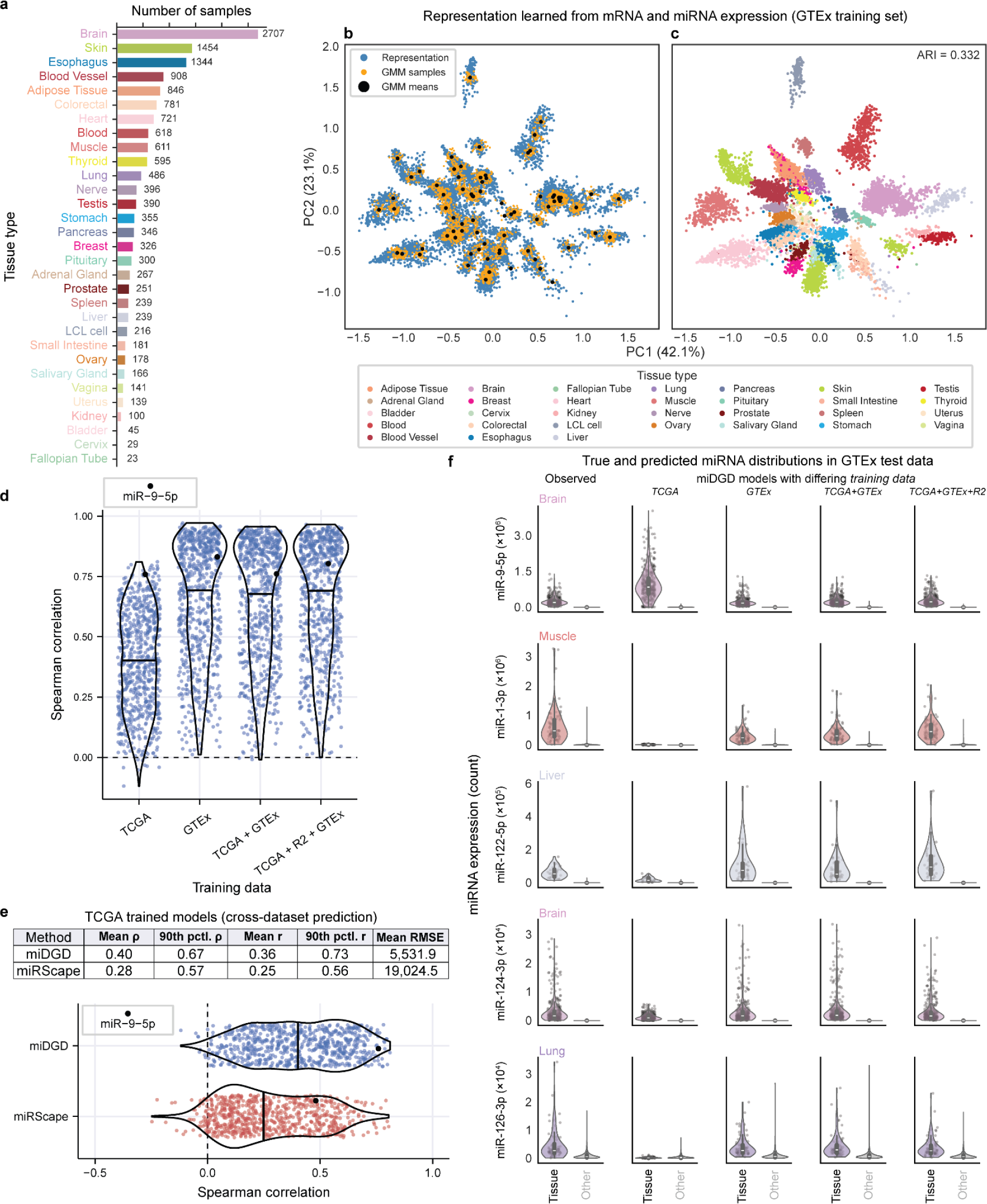
Cross-dataset miRNA prediction in human healthy tissues. **a**, GTEx sample overview, stratified by tissue type. **b**, PCA of latent space for GTEx training samples, showing sample representations (blue) together with GMM samples (orange) and component means (black). **c**, Latent space with sample representations coloured by tissue type. **d**, Violin plots depicting Spearman correlation between cross-tissue miRNA expressions (n = 728) and corresponding predictions based on miDGD models with differing training data. Medians are indicated. “+” indicates the union of training data from independent datasets. **e**, Performance of cross-dataset miRNA expression prediction (top) for supervised methods. Both models are trained on pan-cancer TCGA data while predicting on GTEx test data. The full Spearman correlation distributions are shown (bottom), and the medians are indicated. **f**, Violin plots showing the true expression of selected tissue-specific miRNAs as well as predicted expression from miDGD models trained on various datasets.

When miDGD is trained on GTEx, inferred latent representations cluster in agreement with known tissue labels (ARI = 0.33; Fig. 4b,c), and less strongly when using the TCGA-trained model (ARI = 0.20; Supplementary Fig. 1c,d). Each tissue is described by at least one GMM component, many of which show high tissue-label concordance (Fig. 4b,c; Supplementary Fig. 3a). Accordingly, miDGD achieves reliable cross-tissue miRNA prediction performance (mean ρ≥0. 64 and *r*≥0. 68; Fig. 4d, Supplementary Fig. 4a).

We evaluated cross-dataset generalization by applying TCGA-trained models on tissue samples from the GTEx test set. In this out-of-distribution setting, miDGD improves cross-dataset miRNA prediction relative to the prior supervised method miRSCAPE (mean ρ = 0. 40 and ρ = 0. 28, respectively; Fig. 4e, Supplementary Fig. 4b-d), indicating that the shared latent representation helps stabilize predictions on novel data. The cross-dataset predictions have lower correlation with observed miRNA counts (mean ρ = 0. 4) than those from GTEx-trained miDGD models (mean ρ≥0. 64), underscoring differences in tissue state (cancer and healthy) and potential technical variation between TCGA and GTEx samples (Fig. 4d, Supplementary Fig. 4a).

This difference is also visible in the latent space. Not all GTEx tissues are present in the TCGA data, and tumor samples typically differ in expression from their healthy tissue of origin. In the TCGA-trained miDGD model, GMM components therefore primarily capture cancer-type-specific structure, and GTEx samples are placed within a latent space geometry that only partially reflects healthy tissue variation. Consequently, many GTEx samples are split across multiple components or assigned to components containing mixed tumor types (Supplementary Fig. 3b), whereas the GTEx-trained model yields more tissue-specific components with higher purity (Supplementary Fig. 3a). This discrepancy may explain the performance difference for miRNA prediction in healthy tissues for models trained with and without GTEx data. It highlights the importance of training data matched to the intended prediction setting.

Looking at the top correlating miRNAs, the TCGA-trained model performs well for lowly expressed and tissue-specific miRNAs, while GTEx-containing models perform well for both universally expressed and tissue-specific miRNAs (Supplementary Fig. 5). Examples include well-known cases such as the neuronal miR-9-5p (ρ between 0.76 and 0.83 for the miDGD models) and miR-124-3p (ρ = 0. 54 to 0. 56), the muscle-specific miR-1-3p (ρ = 0. 63 to 0. 90), liver-specific miR-122-5p (ρ = 0. 18 to 0. 33), and lung-specific miR-126-3p (ρ = 0. 36 *to* 0. 93); Fig. 4f, Supplementary Fig. 5).

### Single-cell microRNA Expression Prediction in Human Cell Lines

To explore miRNA prediction at the single-cell level, we evaluated miDGD performance on a whole transcriptome scRNA-seq held-out test set from the Smart-seq-total data (n = 153; Fig. 5a-b, Supplementary Fig. 6a-c)^18^. The Smart-seq-total protocol jointly sequences long and short transcripts, with miRNAs representing a small fraction of the total RNA body and potentially being prone to dropout events^18^. The miRNAs have a mean raw expression count of 0.36, with mir-16 and mir-222 being the most abundant (Fig. 5b).

**Fig. 5:**
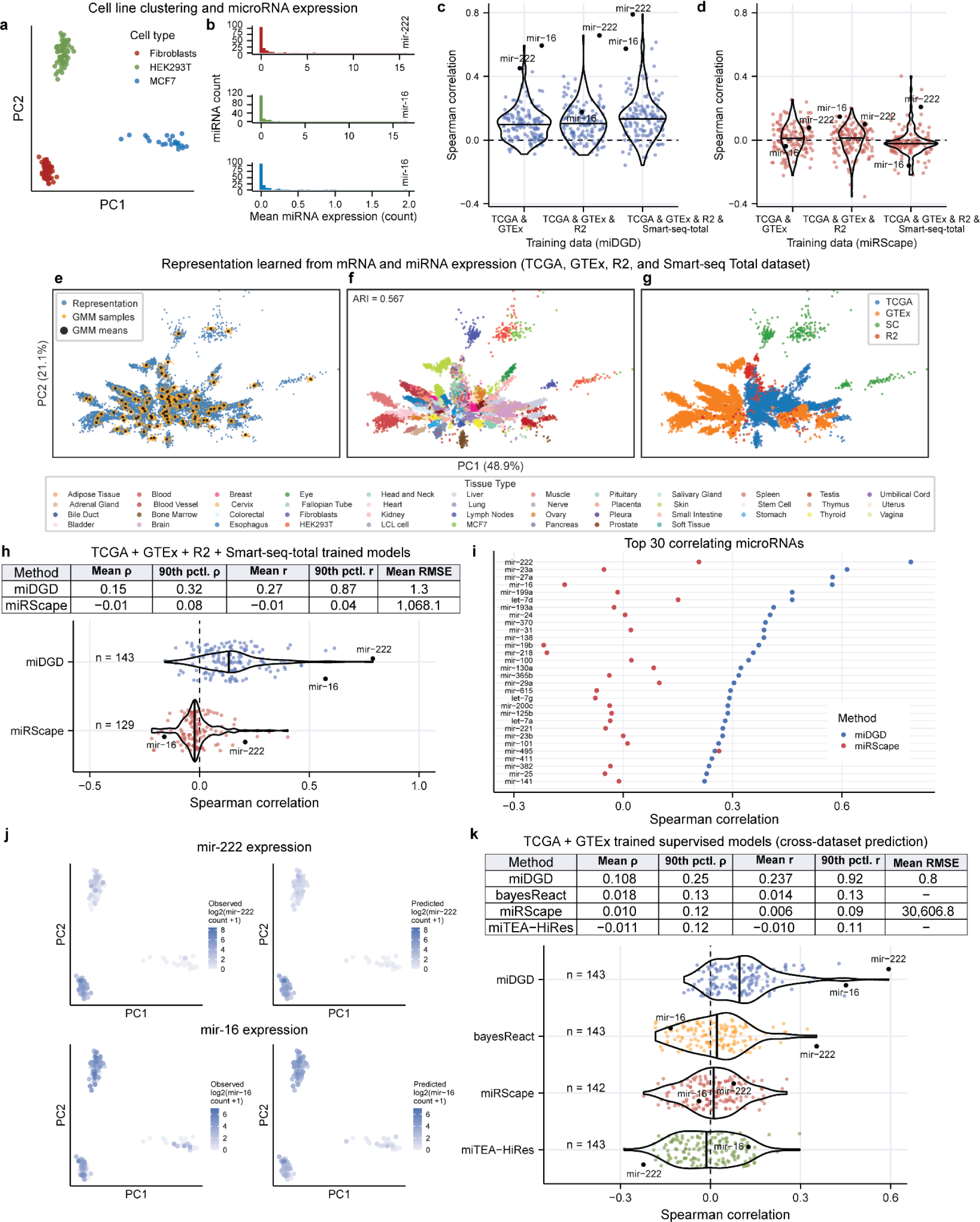
Single-cell miRNA expression prediction and performance. **a**, PCA-based clustering of cells in test data from the Smart-seq-total sequenced human cell lines (n = 153). **b**, Mean miRNA expression (n = 156) is depicted for each cell line, with the most abundant miRNA annotated. **c**, Spearman correlation between observed and predicted expression for miDGD trained on different training data. The median correlation is highlighted, and 143 of 156 miRNAs have sufficient expression variability in the test data to be evaluated. **d**, Spearman correlation for miRSCAPE models with differing training data, matching panel b. miRNAs with minimal expression or prediction variability could not be evaluated, including 14, 13, and 27, respectively. **e**, PCA of latent space for TCGA+GTEx+R2+Smart-seq-total training samples, showing sample representations (blue) together with GMM samples (orange) and component means (black). **f**, Analogous to panel c, showing sample representations colored by tissue type; and **g**, showing sample representations colored by dataset. **h**, Correlation performance for miDGD and miRSCAPE trained on TCGA+GTEx+R2+Smart-seq-total while predicting on the Smart-seq-total test data (top). Spearman correlations between observed and predicted miRNA expression profiles are shown (bottom). pctl, percentile. **i**, Highlight of top 30 miDGD correlating miRNAs from panel e. **j**, Observed (left) and predicted (right) expression across cell clusters for mir-222 (top) and mir-16 (bottom). See panel a for cell line annotations. **k**, Cross-dataset performance for both supervised and unsupervised methods, not relying on within-dataset miRNA expression profiles for training.

We first considered whether a miDGD model trained solely on bulk data could generalize to single cells. Encouragingly, bulk-trained miDGD resulted in positive mean correlations between observed and predicted miRNA expression (n = 143; mean ρ≥0. 11 and *r*≥0. 23; Fig. 5c-d, Supplementary Fig. 6d). Adding single-cell training data improved performance (mean ρ = 0. 15 and ρ = 0. 27; *p* < 0. 05, two-sided Wilcoxon rank-sum test). Hence, training data resembling the test data, e.g., the same cell lines rather than bulk cell mixtures, can refine latent clustering and miRNA predictions (Fig. 5e-g, Supplementary Fig. 6a-c). This extends the coverage of the latent space to regions occupied by single-cell representations. Cells may otherwise be assigned to relatively distant GMM components from the most adjacent bulk samples, with low overlap in representations likely caused by biological and technical differences between immortalized cell lines and primary tumors and tissues^18,48^ (Fig. 5g).

Meanwhile, miRSCAPE produces a lower mean correlation with the single-cell training data included, which can indicate higher susceptibility to technical variation (mean ρ≤0. 01 and *r* < 0. 01 before, and mean ρ =− 0. 01 and *r* =− 0. 01 after inclusion; Fig. 5c-d, Supplementary Fig. 6e,f). This may in part be due to miRSCAPE training independent models for each miRNA, whereas miDGD jointly models the full mRNA and miRNA modalities. Furthermore, performing cross-dataset miRNA predictions, miDGD (mean ρ = 0. 11) outperforms all recent predecessors (mean ρ < 0. 02; Fig. 5k, Supplementary Fig. 6g). Previous methods mainly recover positive correlations for a small number of miRNAs, whereas miDGD extends positive predictive performance to a broader set of miRNAs, including highly abundant instances (Fig. 5).

Top miDGD predictions, in terms of correlation, include the highly abundant mir-16 (ρ = 0. 57) and mir-222 (ρ = 0. 79), as well as several other miRNAs highlighted by Isakova et al.^18^ (mir-27A, several miRNAs from the let-7 family, mir-19b, and mir-25; ρ≥0. 23; Fig. 5h-j).

### miDGD Evaluation on an Independent Single-Cell Test Set

miDGD was further evaluated on an independent test set comprising cells from four human cell lines (A549, 293T, K562, and HeLa; n = 2,310; Fig. 6) to assess performance beyond the datasets used for model training. Contrary to Smart-seq-total sequencing, here, mRNAs and miRNAs were divided into separate paired fractions and sequenced following the PSCSR-seq protocol^19^. This leads to better-resolved miRNA profiles with a mean raw expression count ≥ 9. 55 for each cell line (Fig. 6a).

**Fig. 6:**
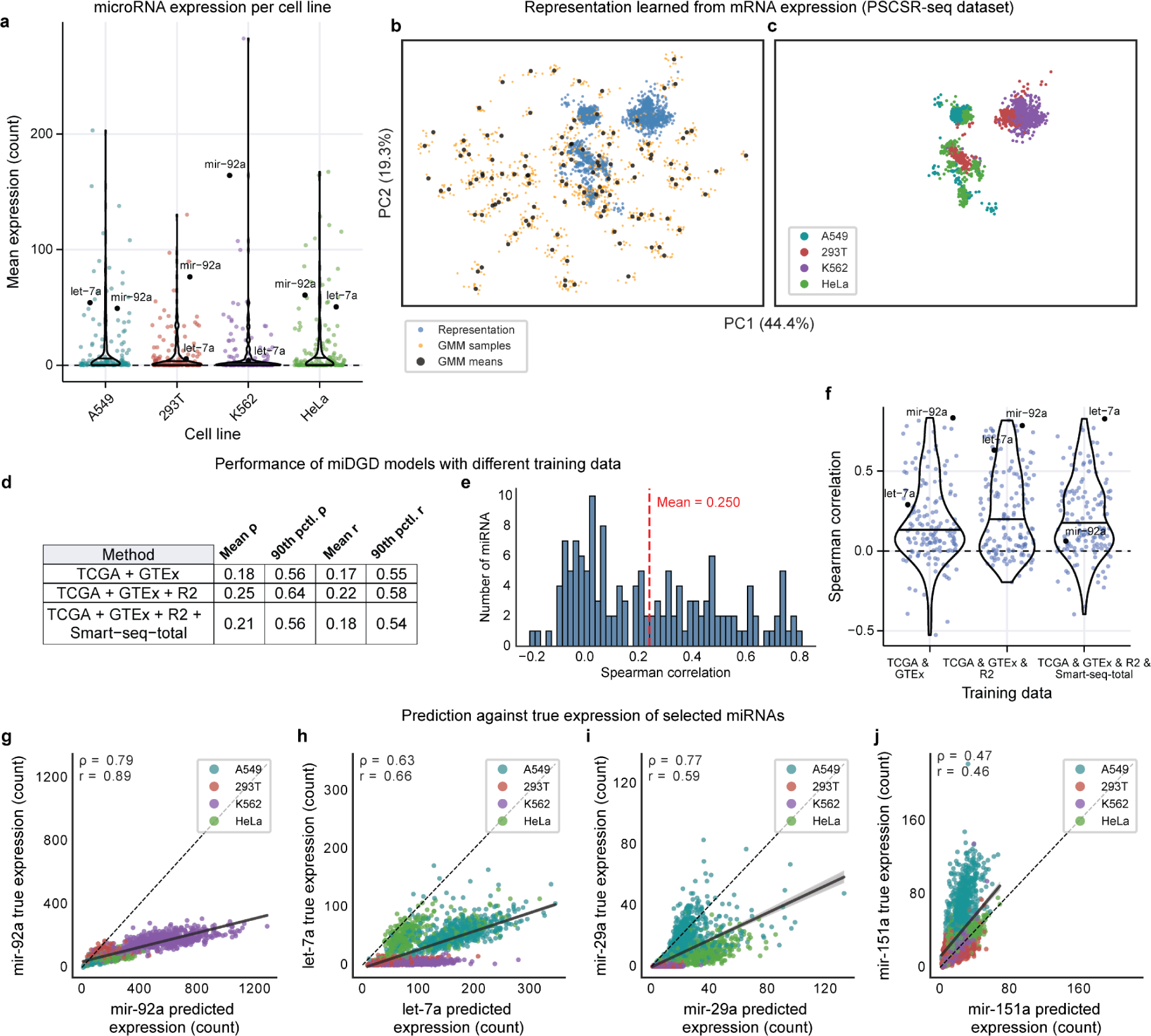
miDGD miRNA single-cell prediction for an independent human cell line dataset. **a**, Mean miRNA expression for each cell line (146 of 156 miRNAs are expressed in the PSCSR-seq human cell line dataset; n = 2,310 cells). The median is highlighted on each violin plot. **b**, Latent space PCA of PSCSR-seq dataset representation (blue) together with GMM means (black) and GMM samples (orange) from TCGA+GTEx+R2 miDGD model. **c**, Analogous with panel b, showing sample representations colored by cell line. **d**, Spearman correlation between observed and predicted test expression for miRNAs (n = 146) in the PSCSR-seq dataset. **e**, Mean performance metrics for different miDGD models, comparing observed and predicted miRNA expression. pctl, percentile. **f**, Spearman correlation between observed miRNA expression and miDGD predictions based on different underlying training data. **g**, True and predicted expression of miR-92a; **h**, let-7a; **i**, miR-29a; and **j**, miR-151a.

By inferring latent representations for the new datapoints under the bulk-trained miDGD model, we observe that some cells locate near multiple GMM component means, while others map to regions of latent space with low support (GMM ARI = 0.004, Fig. 6b-c). Encouragingly, the learned representations cluster cells largely by cell line (K-means ARI = 0.44, see Methods; Fig. 6c), indicating that cell-line structure is present in the latent space even though it is not well captured by the bulk GMM prior.

miRNA predictions from miDGD trained strictly on bulk expression data tend to correlate positively with the observed single-cell miRNA expression (n = 146; mean ρ≥0. 18 and *r*≥0. 17; Fig. 6d-f, Supplementary Fig. 7a-b). Including bulk cell line expression from the R2 RNA Atlas during training improved performance (mean ρ = 0. 25 and *r* = 0. 22; 80% with *r* > 0 and 5.5% with *r* > 0. 75; Fig. 6d-f). However, adding single-cell data from the Smart-seq-total sequenced cell lines (Fig. 5a) did not improve miRNA predictions on average (Fig. 6f; Supplementary Fig. 8; *p* > 0. 05, two-sided Wilcoxon rank-sum test), which was the case for the Smart-seq-total test set (Fig. 5b). This may be due to technical variation from the markedly different sequencing protocols.

miDGD trained on bulk expression data successfully recovered miRNA expression profiles from several abundant miRNAs (Supplementary Fig. 7c), including mir-92a (ρ = 0. 79), highly abundant in K562 cells, and the ubiquitously expressed let-7a (ρ = 0. 63), as well as the lung and A549 specific mir-29a (ρ = 0. 77) and mir-151a (ρ = 0. 47; Fig. 6g-j, Supplementary Fig. 8f-k), several of which Li et al. also highlight^19^. Despite differences between the bulk RNA-seq training set and the zero-inflated scRNA-seq test data, miDGD manages to capture meaningful mRNA-miRNA relationships at the single-cell level.

## DISCUSSION

MicroRNAs represent a critical layer of post-transcriptional gene regulation, yet their abundance is typically unmeasured in single-cell transcriptomic assays. In addition, single-cell miRNA expression prediction remains challenging and poorly solved due to complex regulatory mechanisms acting on mRNAs, as well as detection limitations and technical bias^24,25^. In this study, we introduced miDGD, a deep generative model that learns a shared latent representation of paired mRNA and miRNA expression profiles and enables the prediction of miRNA abundance from mRNA measurements alone. By jointly modelling both modalities within a unified latent space, miDGD captures regulatory structure and provides a novel approach for inferring missing miRNA expression.

Across large bulk datasets from TCGA^34^ and GTEx^35^, we showed that miDGD learns latent representations that reflect major tissue and cancer-type expression patterns. Despite being trained without explicit labels, this organization was preserved when inferring representations from either mRNA or miRNA alone. This indicates that the model captures coherent cross-modal structure rather than overfitting to one data type. More broadly, this extends recent deep generative work showing that multimodal representations can recover biologically meaningful organisation^29,32^. Furthermore, miDGD’s joint modelling can capture cross-modality regulatory relationships, improving robustness and cross-dataset generalization, compared to activity inference approaches^23,24^ that estimate relative regulatory effects from target depletion alone, or the regression-based miRSCAPE^25^, which models miRNAs independently.

We found that miDGD can reconstruct miRNA expression and accurately recovers both tissue-specific and broadly expressed regulators, including miR-9-5p, miR-21-5p, and members of the let-7 family. Predicted miRNA abundances preserved expected relationships with known target genes^41,42^, indicating that reconstructed expression remains consistent with known regulatory interactions^43–45^. Because inference operates in latent space rather than directly on the full gene set, miDGD can also generate miRNA predictions when only a subset of mRNA features is observed. Furthermore, we observed robust performance under strong library size downsampling, suggesting that the learned mapping from mRNA to miRNA remains stable even under sparse input conditions typical of scRNA-seq. Consistent with this, we observed that predictions correlated positively with experimentally measured miRNA levels in independent single-cell datasets, despite the model being trained on bulk data. Prediction accuracy improved when training data more closely matched the biological context of the target dataset, highlighting the importance of diverse, representative training cohorts. However, expanding the training cohort alone did not uniformly improve miRNA predictive performance. In our experiments, incorporating additional datasets, such as the R2 RNA Atlas and single-cell data, did not affect average performance but did slightly reduce prediction accuracy for several prominent miRNAs, suggesting that heterogeneity across biological contexts and sequencing protocols can introduce conflicting signals during training.

miDGD relies on paired mRNA–miRNA training data and is therefore currently limited to species where such datasets are available at a sufficient scale. However, the lack of an encoder allows deep generative decoders to be trained on much smaller datasets than, for instance, variational autoencoders^49^, potentially enabling training on smaller available mouse datasets as well^18,19^. Variability in miRNA annotation and measurement quality may influence predictions, particularly for poorly characterized and lowly expressed transcripts. Many miRNAs in miRBase are likely false positives^50,51^, which we partially mitigate by focusing on the intersection of robustly detected miRNAs across multiple datasets.

The generative architecture underlying miDGD provides multiple opportunities for further development. Because the model learns a shared latent representation, it can, in principle, be trained using partially observed modalities, enabling use on datasets lacking either mRNA or miRNA measurements. The method could also be extended to incorporate additional omics layers such as proteomics and epigenomics, allowing broader multi-omics modelling of gene regulation, which may yield a latent geometry with improved tissue and cell type separation. Moreover, miDGD’s architecture could, in principle, be adapted to other species, as more paired datasets become available, allowing the model to learn conserved regulatory pathways. Explicit modelling of batch and dataset effects within the latent space may further improve cross-cohort robustness. As larger collections of datasets become available, we expect generative modelling approaches such as miDGD to become increasingly powerful tools for reconstructing regulatory information that is not directly measured.

## METHODS

### Data Acquisition and Processing

miDGD was trained and evaluated using paired miRNA and mRNA raw read counts from the same samples or cells. Bulk expression data were acquired from TCGA pan-cancer primary tumor samples^34^. The mRNA counts were obtained from the Recount3 project^52^, and miRNA counts, together with sample annotations, through the GDC data portal^53^. PAM50 and iCluster subtyping were retrieved from TCGAbiolinks^54^. Paired expression data were also retrieved from healthy tissues via the GTEx project^35,47^, and from cell lines through the R2 RNA Atlas^36^. The biological source (tissue of origin) for the cell lines in the R2 RNA Atlas was manually matched to the GTEx tissue labels, ensuring consistent naming and cluster annotation. In bulk datasets, mature miRNA expression is separated into 5p and 3p arms unless otherwise stated.

For single-cell evaluation, we used paired mRNA and miRNA expression data from Isakova et al.^18^ and Li et al.^19^, generated with Smart-seq-total and PSCSR-seq V2, respectively. These datasets comprised HDFa fibroblasts, HEK293T, and MCF7 cells for Smart-seq-total, and A549, 293T, K562, and HeLa cells for PSCSR-seq V2.

In all instances, gene and miRNA annotations were based on GENCODE v.32^18,55^ and miRBase v.22^56^, respectively. The datasets were split into training, validation, and test sets, except for the PSCSR-seq data, which served as an external test set. An overview of the data split and dimensionality is provided in Supplementary Table 1. For models trained on both bulk and single-cell expression data, or used to predict on single-cell test data, we collapsed the 5p and 3p expression counts. Throughout, we use the subset of protein-coding transcripts (mRNAs) and miRNAs detected across multiple bulk datasets (uncollapsed 5p and 3p miRNA expression), as well as the single-cell data (collapsed miRNA arms; Supplementary Table 1).

From the TCGA data, we generated ten downsampled mRNA expression count matrices with varying library sizes for model testing under increasing count sparsity. The matrices were generated by sampling genes with replacement a fixed number of times (1,000 - 10,000,000) and using their sampling counts as expression values. Each gene was sampled with probability proportional to its normalized read counts per kilobase of transcript per million mapped reads (RPKM).

### Model Architecture

The miDGD model is based on the multiDGD model and introduces several modifications that are optimized for miRNA expression prediction from mRNA expression data^32^, including modality-specific decoder branches, a shared latent space for both modalities, and distribution choices adapted to sparse miRNA counts.

miDGD represents each sample *i* as a low-dimensional latent vector *z_i_*, shared across both the mRNA and miRNA modalities. The latent space is modelled by a Gaussian mixture model (GMM) with parameters **φ** (mixture weights, component means, and diagonal covariance matrices) and *K* the number of mixture components. Together, *Z* and **φ** define the latent variables and their distribution, allowing the model to capture heterogeneous biological states in a compact space and to organize samples across cancer and tissue types.

For each sample *i*, the latent vector *z_i_* has length *L*, and the decoder θ, implemented as a fully connected network, maps this shared representation into modality-specific predicted mean counts for mRNA and miRNA through two output branches (Fig. 1c). These predicted mean counts are scaled by the sample-specific library sizes (sum of mRNA or miRNA expression counts in that sample) and used as the mean parameters of the negative binomial and zero-inflated negative binomial distributions that model mRNA and miRNA counts, respectively. All parameters (decoder weights, GMM parameters, and sample-specific representations) are updated jointly by backpropagation during training. The latent space dimension *L* and the number of mixture components *K* were treated as tunable hyperparameters.

### Model Training

During training, miDGD optimizes a joint objective that combines reconstruction of both modalities (mRNA and miRNA) with a Gaussian mixture model (GMM) prior over the latent representations (Fig. 1c). The objective is to minimize the loss function:

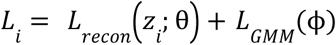

The decoder maps the latent representation *z_i_* to the parameters of the count distributions. The reconstruction term is defined as the sum of negative log-likelihoods under a negative binomial (NB) for mRNA and a zero-inflated negative binomial (ZINB) for miRNA. The NB distribution models overdispersed count data typical of RNA-seq, whereas the ZINB additionally captures the excess zeros characteristic of sparse small RNA-seq measurements. Both are parameterized by decoder outputs and scaled by sample-wise library size. The reconstruction loss is defined as:

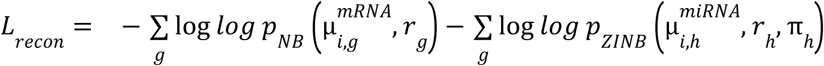

where 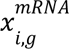 and 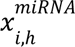 denote the observed mRNA and miRNA counts for sample *i*, gene *g* and miRNA ℎ, respectively; 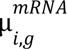 and 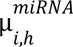 are the corresponding decoder-predicted mean counts after scaling by the sample-specific library size; *r_g_* and *r*_ℎ_ are feature-specific dispersion parameters for mRNAs/genes and miRNAs; and µ_ℎ_ are zero-inflation probabilities for miRNAs.

The GMM prior acts as a regularization term on *z_i_*, given by the negative log-density under the mixture:

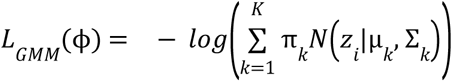

implemented using a numerically stable “log-sum-exp” computation, which evaluates the log of the sum of exponentials of component log-densities while avoiding numerical underflow and overflow, with current means µ*_k_*, diagonal covariances Σ*_k_* (stored as log-variances) and weights π*_k_*.

The loss is averaged across features and samples, and gradients are backpropagated through the decoder, the GMM parameters, and the sample-specific representations. The decoder and GMM parameters are updated per mini-batch, whereas sample-specific latent representations are updated once per epoch, using the AdamW optimizer.

### Hyperparameter Tuning

Hyperparameters were optimized using the Weights & Biases (wandb) sweeps framework^57^ that enabled systematic exploration of architectural and training choices for all miDGD models across dataset combinations. For each candidate setting, a complete training run was performed with a fixed train-validation split, and the hyperparameter selection criterion was the best miRNA reconstruction loss. This procedure was repeated for each model-dataset combination, ensuring that the final model choices reflected the specific properties of the training data (e.g., TCGA-only or multiple-dataset settings) rather than a single global configuration. Subsequently, the optimal parameters for each setting are used.

### Training Dataset Configuration

The complete set of models, with varying combinations of training and validation data, is: TCGA-only, GTEx-only, TCGA+GTEx, TCGA+GTEx with collapsed 5p and 3p miRNA expression, TCGA+GTEx+R2, TCGA+GTEx+R2 with collapsed 5p and 3p miRNA expression, and TCGA+GTEx+R2+Smart-seq-total with collapsed miRNA. Here, “+” denotes the union of training datasets as well as validation sets. Optimal hyperparameters for each model are reported in Supplementary Table 3.

### Inference on New Samples

Prediction of miRNA expression for new samples was performed in two stages using the trained miDGD model, with only mRNA expression provided as input.

Given a new dataset *X_test_* containing only mRNA expression profiles, each sample was first assigned an initial latent representation by exploiting the learned Gaussian mixture over the latent representation. Specifically, the means of the fitted GMM components were used as a set of representation candidates, and each candidate was decoded to predict mRNA expression. The negative log-likelihood under the negative binomial distribution output model is computed with respect to the observed true mRNA expression. For each sample, the GMM mean that yielded the lowest mRNA reconstruction loss was selected as the initial representation.

In the second step, these initial latent representations were refined by gradient-based optimization of *z_i_*, with all decoder and GMM parameters held fixed at their trained values. For a predefined number of optimization steps, each representation was updated to minimize the sum of the mRNA reconstruction loss and the GMM loss. The final latent representation was then passed through the decoder to obtain predictions for both mRNA and miRNA expression. In the output layer, decoder logits are transformed with a softmax to yield feature-wise proportions, which are subsequently scaled by a library size to obtain predicted counts.

### Performance Evaluation

Prediction performance was assessed using standard correlation metrics. For each model and dataset, we quantified agreement between observed and predicted miRNA expression using Spearman rank correlation and Pearson correlation, computed per miRNA across samples. Clustering quality of the latent representation was evaluated with the Adjusted Rand Index (ARI) between sample-to-component assignments and known tissue or cancer type labels. For GMM-based clustering, ARI was computed from sample-to-component assignments given by the Gaussian mixture prior, whereas for K-means clustering we applied K-means directly to the latent representations and computed ARI using the resulting cluster labels.

### Sample-to-Component GMM-Based Clustering

After training, we used the fitted GMM to assign each sample to a latent component. For a given representation *z_i_*, we computed the posterior mixture responsibilities γ*_ik_* ∝ π *N_k_* (µ*_k_*, Σ*_k_*) and assigned sample *z_i_* to the component with highest posterior probability, 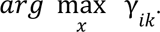. Component purity for a given tissue or cancer type was computed as the fraction of samples of that label assigned to the dominant component, with higher values indicating more tissue-specific placement in latent space.

### Cancer Type Prediction from Latent Components

To evaluate whether the learned latent representation captures cancer-type identity, we used the fitted GMM components as unsupervised predictors of cancer type. For each component, the majority cancer type was determined from training samples assigned to that component. For a new sample, the predicted cancer type was the majority label of the component to which the sample was assigned by maximum posterior responsibility.

Prediction performance was quantified using majority-vote accuracy, defined as the fraction of samples whose predicted cancer type matches the true label. We additionally evaluated the prediction using ARI, homogeneity, and completeness. Homogeneity measures whether each component contains samples of a single cancer type; completeness measures whether all samples of a given cancer type are assigned to the same component. Both metrics range from 0 to 1, with higher values indicating stronger agreement.

### Method Comparisons

We compared miDGD to miRSCAPE (XGBoost)^25,26^, bayesReact^24^, and miTEA-HiRes^23^. miRSCAPE was trained on the same training data as miDGD, and the hyperparameters were defined based on the grid search hyperparameter tuning performed by Olgun et al. on a subset of the TCGA samples^25^. The miRSCAPE wrapper script for XGBoost was updated to ensure all miRNA models were saved for multiple-dataset predictions.

The activity inference methods, bayesReact and miTEA-HiRes, require no prior model training (unsupervised with respect to miRNA abundance) and were applied directly to each full dataset with default settings. Both methods perform activity inference for mature miRNAs separated by the 5p and 3p arms. Subsequently, a primary mature miRNA activity profile was assigned to each collapsed single-cell miRNA expression profile, based on the arm with the highest mean expression across the TCGA and GTEx data.

## ACKNOWLEDGEMENTS

Most of the computing for this project was performed on the GenomeDK cluster. We would like to thank GenomeDK and Aarhus University for providing computational resources and support that contributed to these research results.

We also acknowledge feedback from the Danish Single-Cell Examination Platform (CellX) and the Single-Cell and Spatial Core Facility, Department of Molecular Medicine (MOMA), Aarhus University Hospital, Denmark.

## AUTHOR CONTRIBUTIONS

J.S.P. contributed to conceptualization, supervision, and manuscript review and editing. V.S. and A.K. contributed to methodology through theoretical discussions, expertise in deep generative decoder models, and the initial model implementation. F.Z. contributed to software development by implementing miDGD, conducting hyperparameter optimization, training all miDGD models, and generating predictions across all test sets; M.D. contributed to the early stages of model development. A.M.R. contributed to data curation by retrieving and processing the datasets, with contributions from F.Z. F.Z. and A.M.R. jointly conducted formal analysis, validated and evaluated model performance, prepared visualizations, and wrote the original manuscript draft under the supervision of J.S.P. J.S.P., A.K., V.S., and M.D. contributed to manuscript review and editing.

## DATA AVAILABILITY

All data used during this study are publicly available. Paired mRNA and miRNA expression profiles were obtained from TCGA using the Recount3 project and GDC data portals^34,52,53^, and from GTEx using the GTEx portal (https://www.gtexportal.org/home/).

Meanwhile, cell line and single-cell expression data were obtained from the Gene Expression Omnibus (GEO) through the accession numbers GSE138734 (R2 RNA Atlas), GSE151334 (Smart-seq-total), and GSE226714 (PSCSR-seq). R2 RNA Atlas annotation files were retrieved from the R2 platform (https://r2platform.com/rna_atlas/).

The miDGD model and code are available at https://doi.org/10.5281/zenodo.21886227. Jupyter notebooks and scripts to reproduce our analysis and figures are available at https://doi.org/10.5281/zenodo.21886206. The dataset is available at https://doi.org/10.5281/zenodo.21885644. The latest release is available at GitHub ( https://github.com/JakobSkouPedersenLab/miDGD and https://github.com/JakobSkouPedersenLab/miDGD_paper).

