## Supplementary for "A deep generative decoder predicts microRNA expression from bulk and single-cell mRNA profiles"

### Supplementary Data

**Table of content**

Supplementary Figure 1
Supplementary Figure 2
Supplementary Figure 3
Supplementary Figure 4
Supplementary Figure 5
Supplementary Figure 6
Supplementary Figure 8
Supplementary Figure 9
Supplementary Figure 10
Supplementary Figure 11
Supplementary Table 1
Supplementary Table 2
Supplementary Table 3

#### Supplementary Figure 1

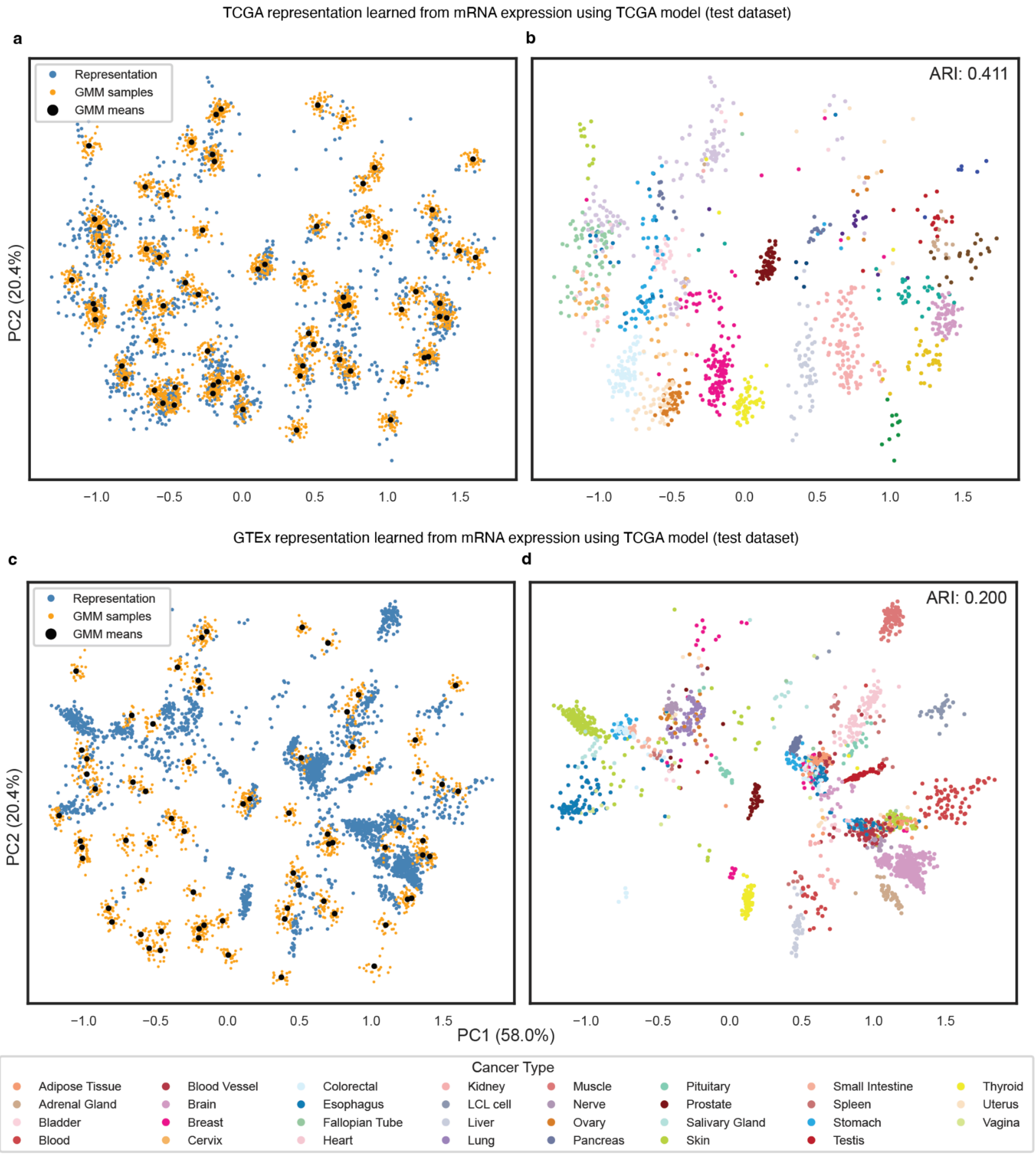


**Supplementary Fig. 1: Latent space structure and GMM components for TCGA and GTEx test dataset.**
**a**, PCA of latent space for TCGA test samples, showing sample representations (blue) together with GMM samples (orange) and component means (black). **b**, Analogous with panel a, showing sample representations colored by cancer type. **c**, Similar to panel a, but using GTEx

test dataset and the representation is optimized using model trained on TCGA dataset. The PCA representation is projected into the same coordinate system as panel a. **d**, Analogous with panel c, showing sample representations colored by cancer type.

#### Supplementary Figure 2

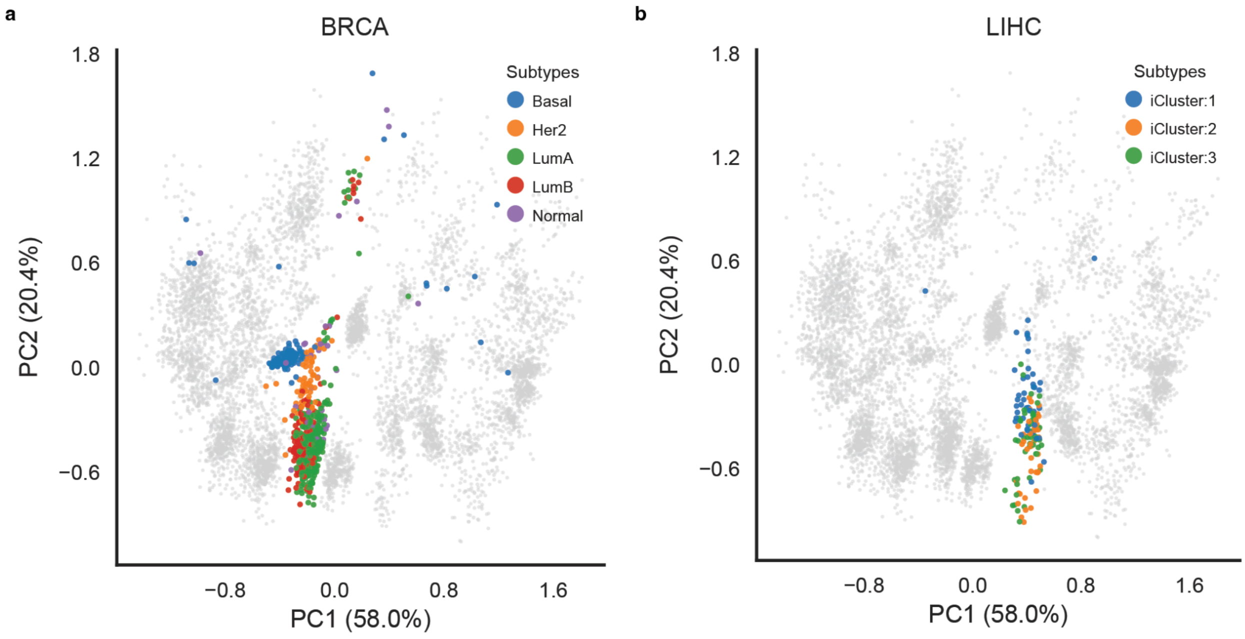


**Supplementary Fig. 2: Latent space visualization of miDGD representations colored by TCGA molecular subtypes.**

**a,** BRCA samples annotated by PAM50 subtypes (Basal, Her2, LumA, LumB, Normal) projected onto the first two principal components of the latent space; remaining samples are shown in grey. **b,** LIHC samples annotated by iCluster subtypes (1–3) on the same latent space, with other samples in grey.

### Supplementary Figure 3

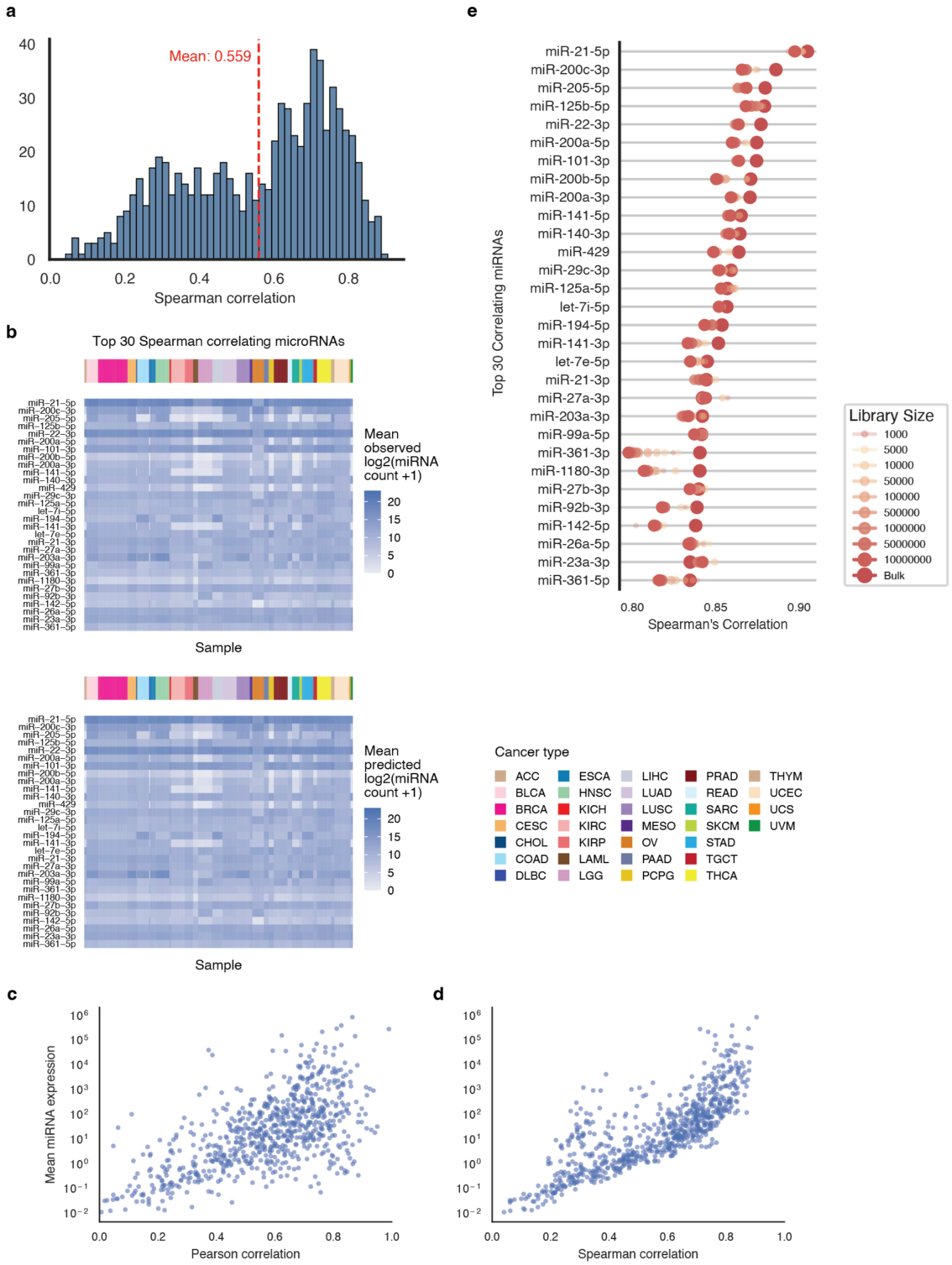

**Supplementary Fig. 3: miDGD predictions across pan-cancer TCGA primary tumor samples.**

**a**, Spearman correlation between observed and predicted test expression for miRNAs (n = 755). **b**, Heatmaps for the mean cancer-type expression (top) and predictions (bottom) for the top 30 Spearman correlating miRNAs. **c,** Pearson correlation and, **d,** Spearman correlation between observed and predicted miRNA expression across samples (x axis) versus mean observed miRNA expression (y axis, log scale), with each point representing one miRNA. **e**, Spearman correlations of the 30 best-predicted miRNAs in the TCGA test data and corresponding semi-synthetic datasets with downsampled gene expression to fixed library sizes.

#### Supplementary Figure 4

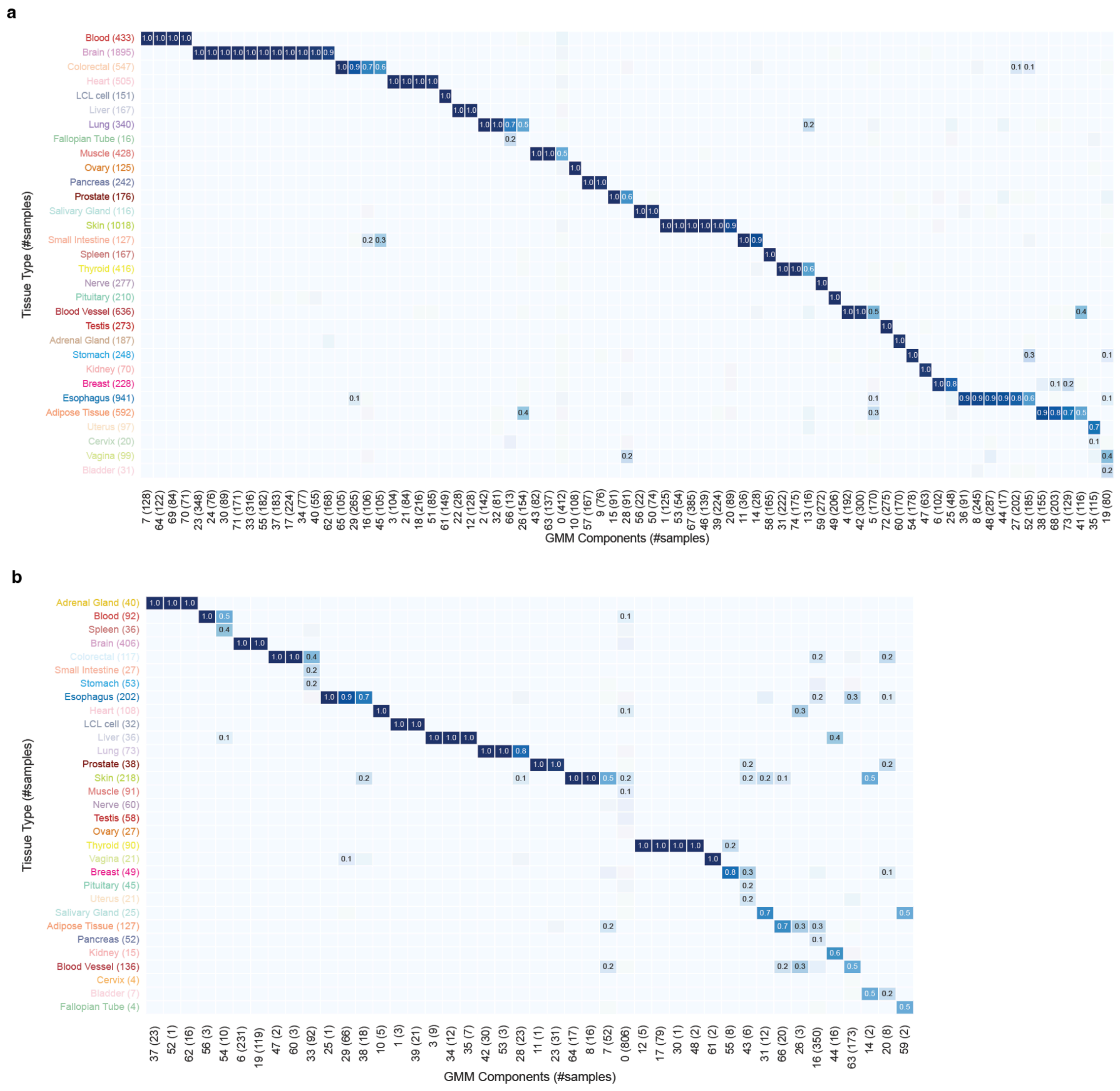


**Supplementary Fig. 4: Tissue-type composition of Gaussian mixture components in GTEx.**
**a,** Matrix showing, for each GTEx tissue type, the fraction of GTEx training samples assigned to each Gaussian mixture component when applying the GTEx-trained model (columns sum to 1).
**b,** Similar to panel a, depicting the GTEx test samples GMM assignment under the TCGA-trained model (columns sum to 1).

**Supplementary Figure 5**

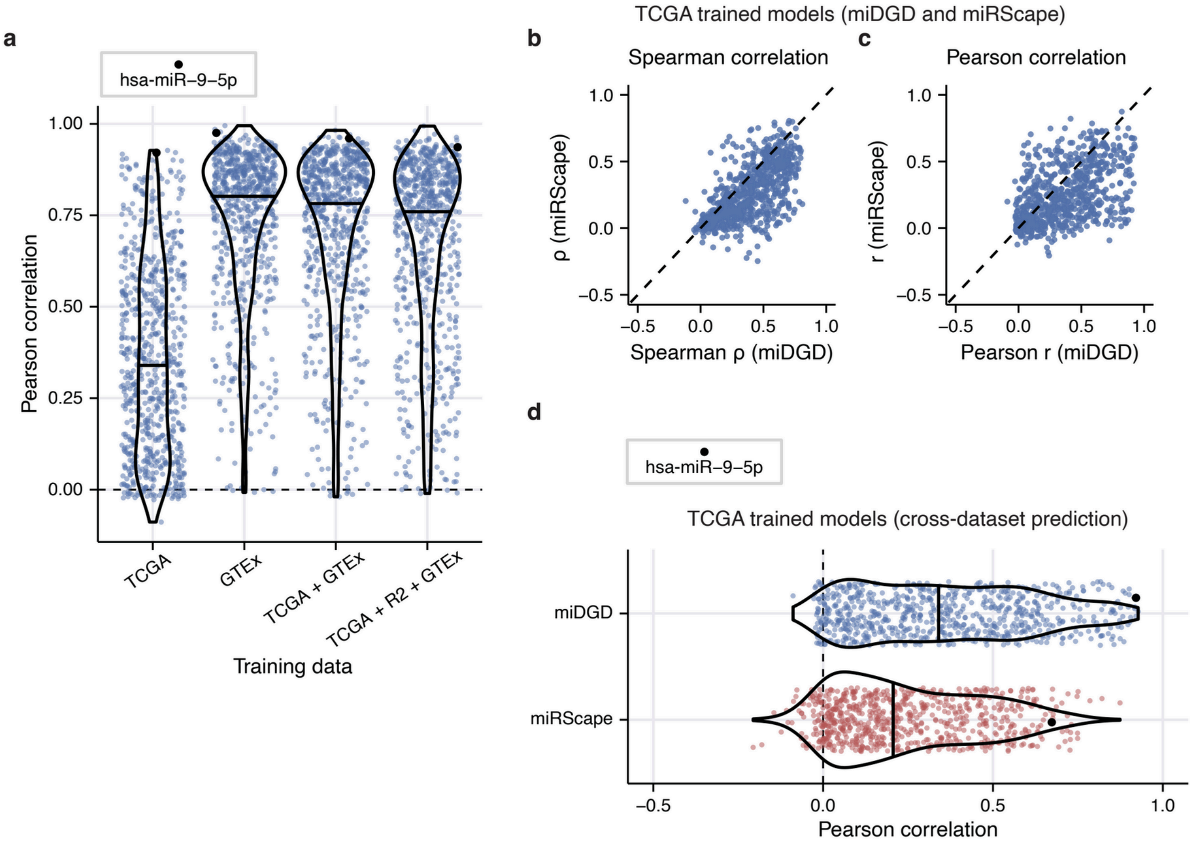


**Supplementary Fig. 5: Extended miDGD and miRSCAPE comparison on GTEx test data.** **a**, Pearson correlations for the GTEx test data between observed and predicted miRNA expression profiles (n = 728), comparing miDGD models with different training sets. Medians are indicated. **b**, Spearman correlation between observed and predicted miRNA expression (n = 728) across the GTEx test data (n = 2,310). The scatterplot depicts correlation coefficients obtained from miDGD against miRSCAPE, and the central line (dashed) is highlighted. **c**, Scatterplot of Pearson correlation obtained from miDGD and miRSCAPE, matching panel b. **d**, Comparison of miDGD and miRSCAPE trained on the TCGA training set and predicting on the GTEx test set. Pearson correlations between observed and predicted miRNA expressions (n = 728) are depicted and median highlighted.

#### Supplementary Figure 6

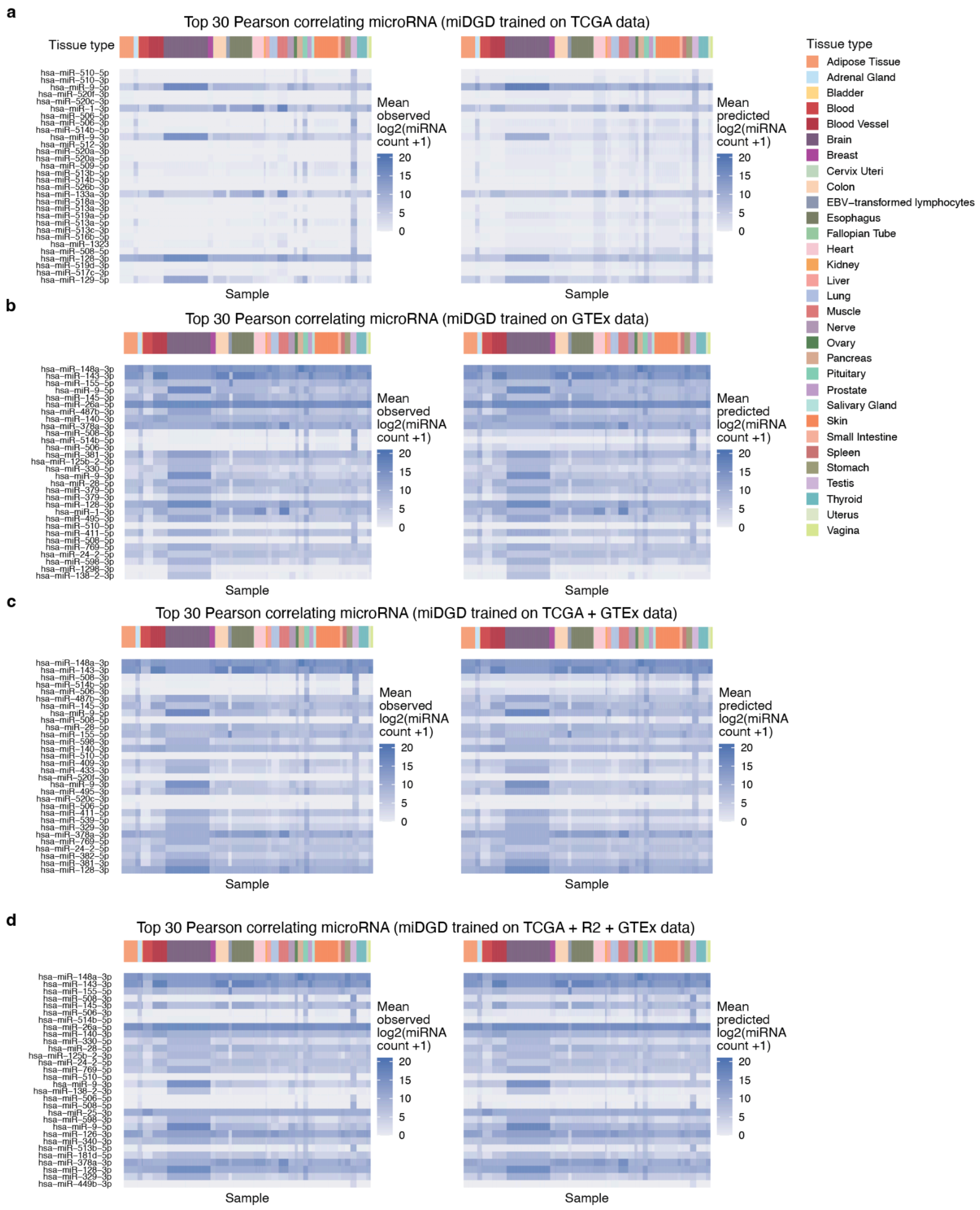


**Supplementary Fig. 6: Top 30 correlating miRNAs from different miDGD models.** Observed (left) and predicted (right) cross-tissue miRNA expression profiles for miDGD models trained on (**a**) TCGA; (**b**) GTEx; (**c**) TCGA+GTEx; and (**d**) TCGA+GTEx+R2 data. Top 30 correlating miRNAs between expression and predictions are shown for each model across the GTEx test samples (n = 2,310).

#### Supplementary Figure 7

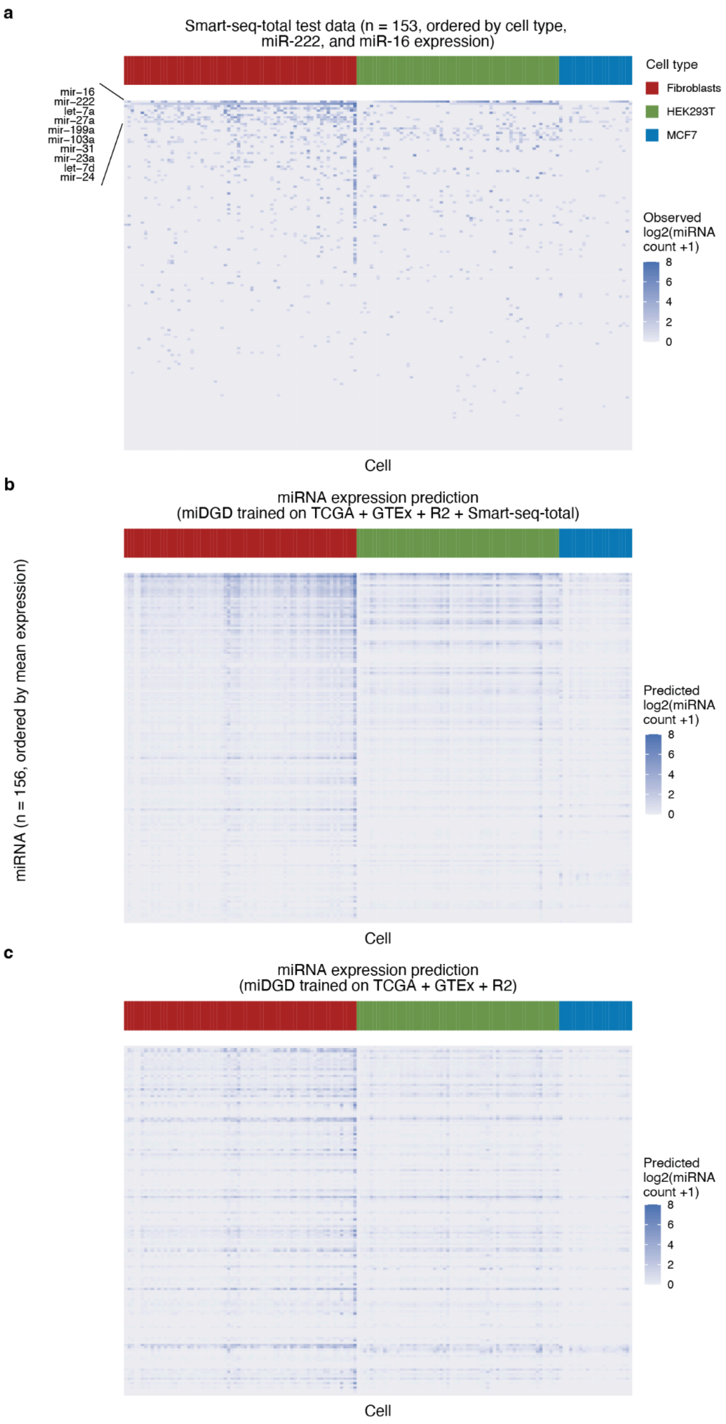


**Supplementary Fig. 7: Observed and miDGD predicted single-cell miRNA expression profiles.**

**a**, Observed expression profiles of all miRNAs (n = 156) across the full Smart-seq-total test set (n = 153). miRNAs with the largest mean expression are annotated (n = 10). **b**, miRNA expression predictions using miDGD trained on the combined TCGA+GTEx+R2+Smart-seq-total training data. Entries match panel a ordering. **c**, Predictions using miDGD trained on TCGA+GTEx+R2 samples.

**Supplementary Figure 8**

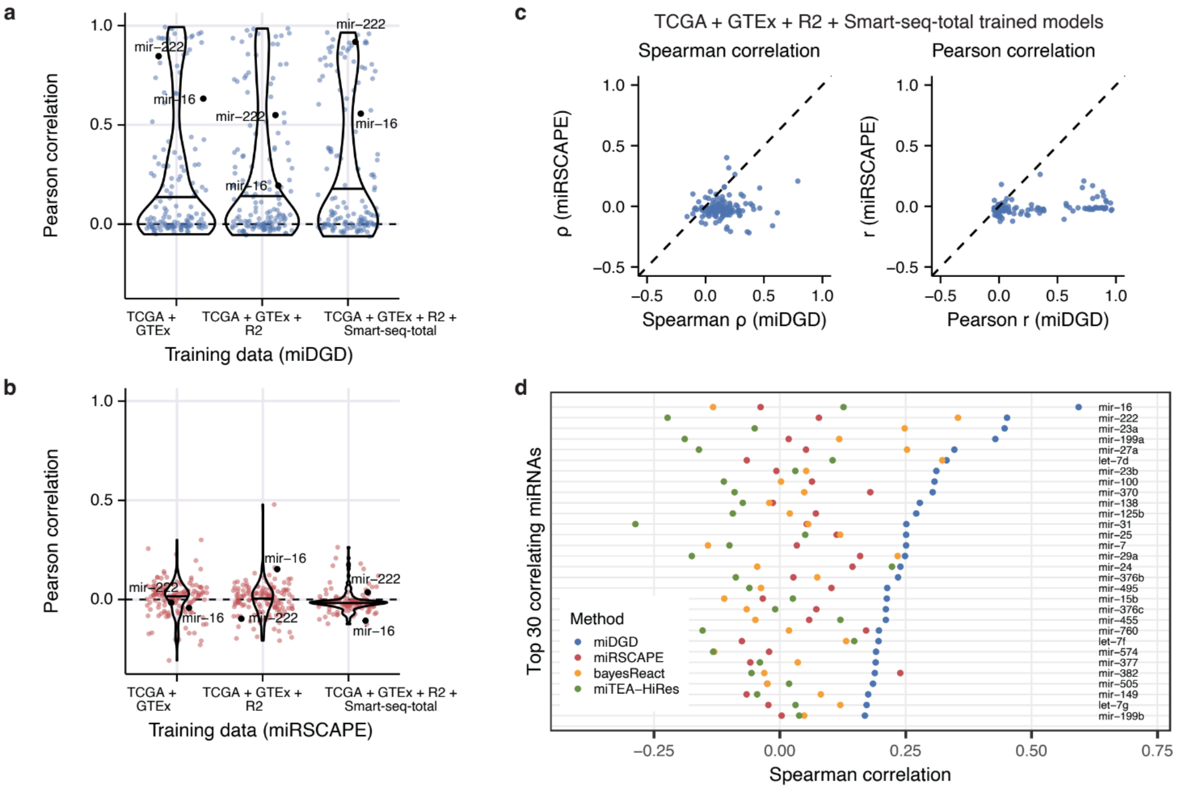


**Supplementary Fig. 8: Method comparisons for single-cell Smart-seq-total test data.**
**a**, Distribution of Pearson correlations between observed and predicted miRNA expression (n = 143) for miDGD models. The median correlation is highlighted. **b**, Pearson correlation for miRSCAPE models with differing training data, and median highlighted similar to panel a. miRNAs with sufficient expression and prediction variability are included, entailing 142, 143, and 129 miRNAs, respectively, for the different models. **c**, miDGD plotted against miRSCAPE performance. Depicted are Spearman correlation (left) and Pearson correlation (right) between observed and predicted miRNA expressions (n = 129). **d**, Top 30 miRNAs based on expression and miDGD prediction correlation from Fig. 5k.

**Supplementary Figure 9**

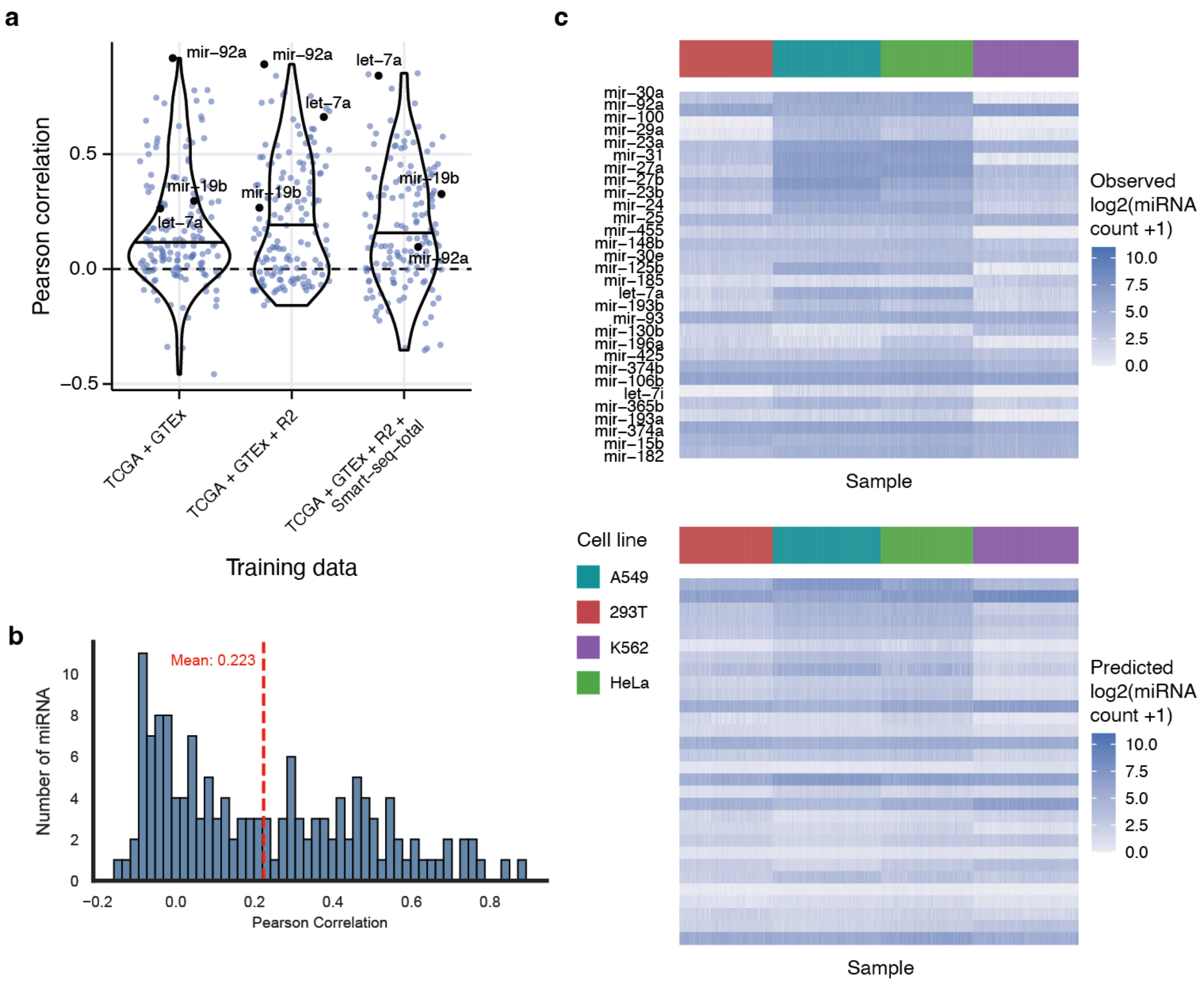


**Supplementary Fig. 9: Single-cell miDGD predictions for PSCSR-sequenced human cell lines.**

**a**, Pearson correlation between observed and predicted miRNA expression across 2,310 cells (n = 146 miRNAs). Predictions are obtained from miDGD models with differing training sets. Median values are indicated. **b**, Pearson correlation distribution for miDGD trained on bulk RNA-seq data (TCGA+GTEx+R2). **c**, Expression profiles observed (top) and predicted (bottom) for top 30 miRNAs with the highest Spearman correlation. Predictions are obtained from miDGD trained on bulk TCGA, GTEx, and R2 RNA Atlas training data.

#### Supplementary Figure 10

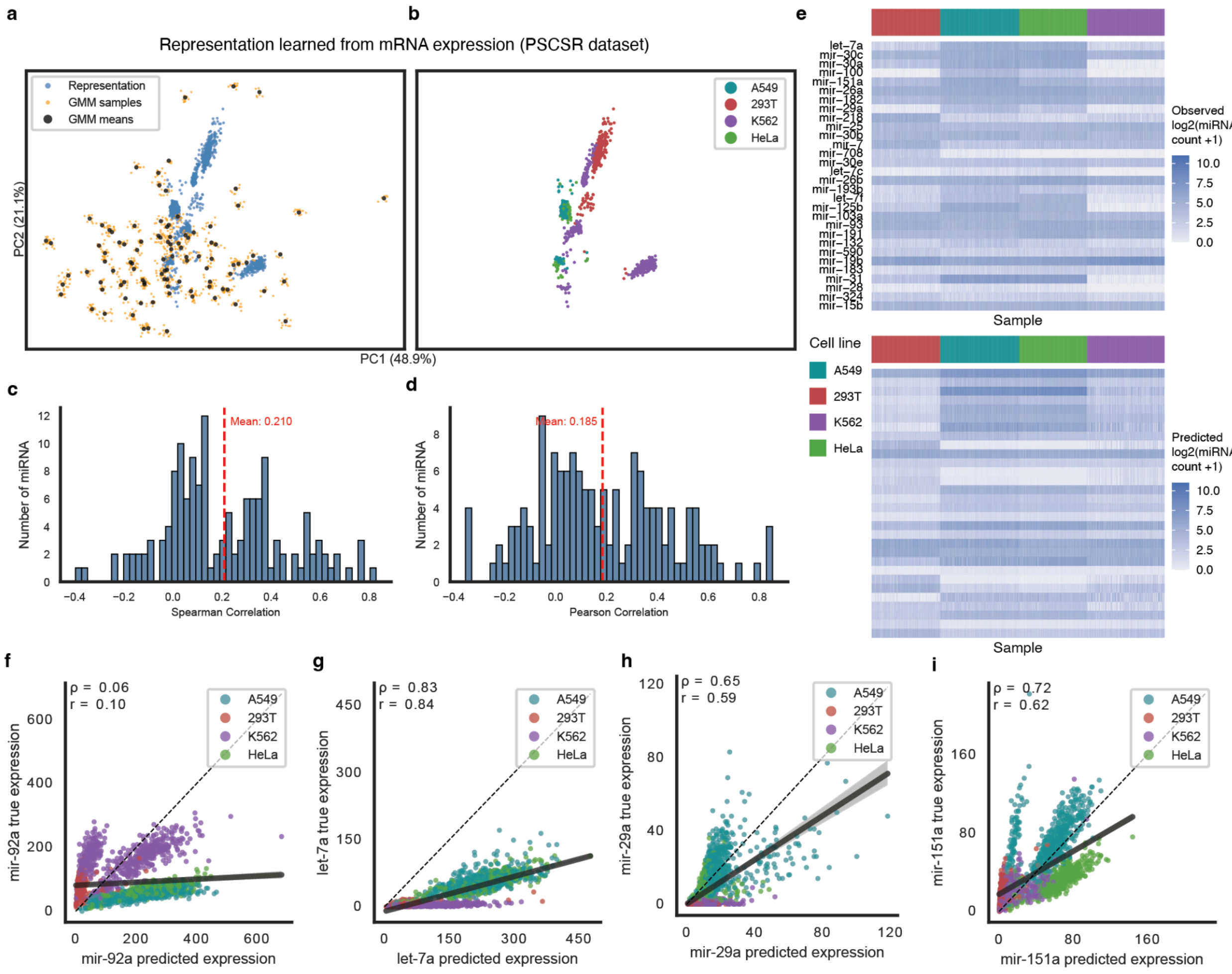


**Supplementary Fig. 10: Single-cell miDGD predictions for PSCSR-sequenced human cell lines using the TCGA+GTEx+R2+Smart-seq-total trained miDGD model.**
**a**, Latent space PCA of PSCSR dataset representation (blue) together with GMM means (black) and GMM samples (orange) from TCGA+GTEx+R2+Smart-seq-total miDGD model. **b**, Analogous with panel a, showing sample representations colored by cell line. **c**, Spearman and, **d**, Pearson correlation between observed and predicted test expression for miRNAs (n = 146). **e**, Observed (top) and predicted (bottom) expression for top 30 Spearman correlating miRNAs. **f**, True and predicted expression of miR-92a; **g**, let-7a; **h**, miR-29a; and **i**, miR-151a.

**Supplementary Figure 11**

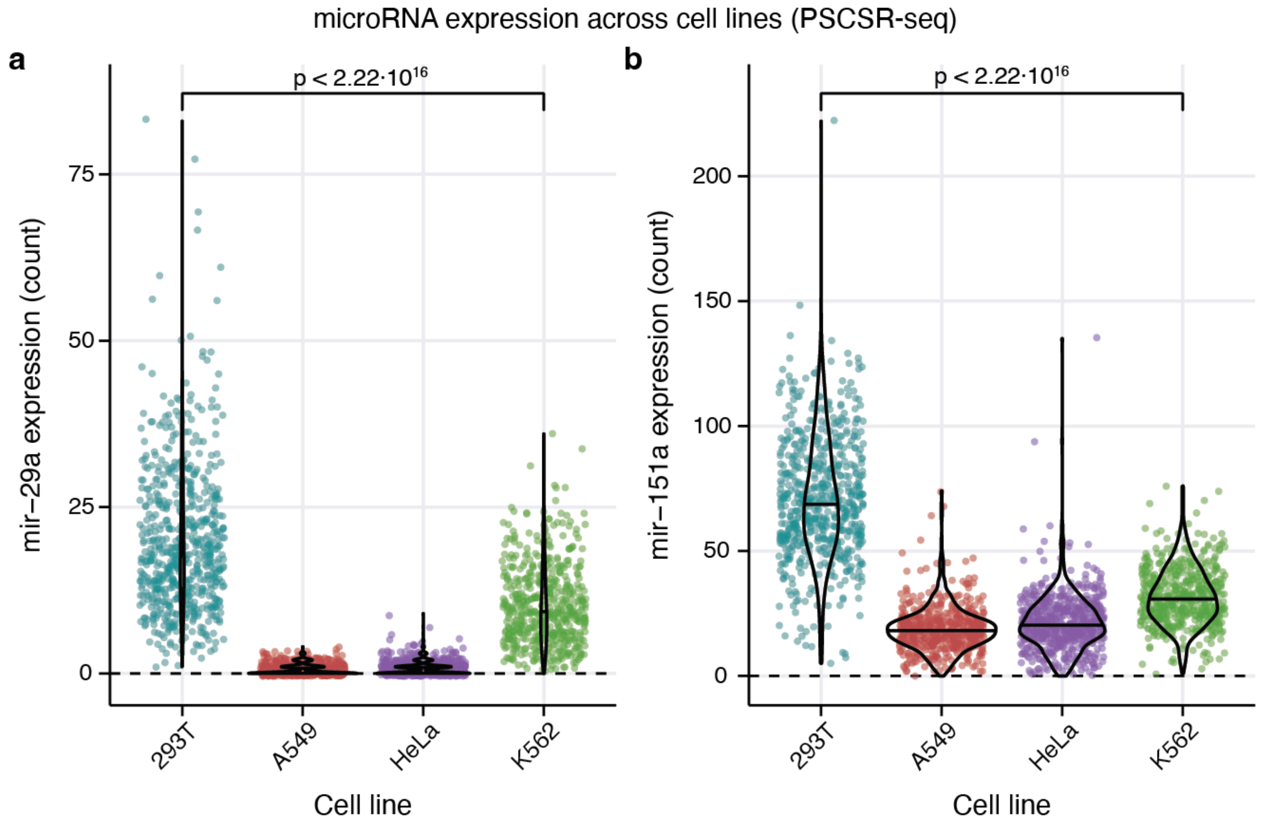


**Supplementary Fig. 11: mir-29a and mir-151a expression across human cell lines (PSCSR-seq).**

**a**, Single-cell mir-29a expression count distribution across four human cell lines (n = 2,310 cells). Median values are indicated on the violin plots. P-value for two-sided Wilcoxon rank-sum test between the two cell lines with the highest median expression is shown. **b**, Similar to panel a, expression distribution for mir-151a.

**Supplementary Table 1**

| Model | Dataset | miRNA 5p/3p | Data dimensions |
|---|---|---|---|
| **1** | TCGA<br>train:val:test split = 75:12.5:12.5 | non-collapsed | Samples: 9640<br>mRNA: 18393<br>miRNA: 755 |
| **2** | GTEx<br>train:val:test split = 70:15:15 | non-collapsed | Samples: 15398<br>mRNA: 18393<br>miRNA: 755 |
| **3a** | TCGA+GTEx | non-collapsed | Samples: 25038<br>mRNA: 18393<br>miRNA: 755 |
| **3b** | TCGA+GTEx | collapsed | Samples: 25038<br>mRNA: 18393<br>miRNA: 418 |
| **4a** | TCGA+GTEx+R2<br><br>R2 samples: 294 (all used for training) | non-collapsed | Samples: 25332<br>mRNA: 18076<br>miRNA: 728 |
| **4b** | TCGA+GTEx+R2<br><br>R2 samples: 294 (all used for training) | collapsed | Samples: 25332<br>mRNA: 18076<br>miRNA: 414 |
| **5** | TCGA+GTEx+R2+Smart-seq-total<br><br>Smart-seq-total cells: 610<br>train:val:test split = 60:15:25 | collapsed | Samples: 25942<br>mRNA: 15625<br>miRNA: 156 |
| **Independent test dataset** | | | |
| | PSCSR-seq | collapsed | Samples: 2310<br>mRNA: 17300<br>miRNA: 418 |

**Supplementary Table 1: Dataset dimensionality and model configuration.**
For each miDGD model, the table lists the dataset combination, train:validation:test split (where applicable), mature miRNA arms (non-collapsed, with 5p/3p arms distinguished, or collapsed, with 5p/3p arms combined), and the number of samples, mRNAs, and miRNAs included in total. A given dataset is split once, and the same split is reused across all models that include it, ensuring the same samples are used for training. In combined models (model 3-5), datasets are concatenated at the sample level and share a common set of genes and miRNAs defined as the intersection across included datasets.

**Supplementary Table 2**

| True Cancer Type | N Samples | Acc | Predicted Cancer Type (misclassified) |
|---|---|---|---|
| ESCA | 23 | 0.000 | STAD (47.8%), LUSC (30.4%), HNSC (17.4%) |
| KICH | 8 | 0.000 | KIRC (100.0%) |
| READ | 20 | 0.000 | COAD (95.0%), UCEC (5.0%) |
| ACC | 10 | 0.200 | PCPG (80.0%) |
| CESC | 38 | 0.395 | HNSC (34.2%), LUSC (7.9%), BLCA (5.3%) |
| LUSC | 59 | 0.559 | HNSC (20.3%), STAD (8.5%), BRCA (6.8%) |
| UCS | 7 | 0.571 | TGCT (28.6%), CESC (14.3%) |
| BLCA | 51 | 0.588 | CESC (11.8%), HNSC (11.8%), LUSC (7.8%) |
| MESO | 11 | 0.636 | SARC (18.2%), PAAD (9.1%), TGCT (9.1%) |
| UCEC | 66 | 0.712 | OV (13.6%), CESC (6.1%), BRCA (3.0%) |
| KIRP | 37 | 0.757 | KIRC (18.9%), BRCA (5.4%) |
| HNSC | 62 | 0.758 | LUSC (21.0%), BRCA (3.2%) |
| LUAD | 63 | 0.762 | STAD (14.3%), PAAD (6.3%), OV (1.6%) |
| LIHC | 46 | 0.826 | KIRC (15.2%), CHOL (2.2%) |
| DLBC | 6 | 0.833 | BRCA (16.7%) |
| STAD | 51 | 0.863 | PAAD (7.8%), CESC (3.9%), COAD (2.0%) |
| PAAD | 22 | 0.864 | SARC (4.5%), STAD (4.5%), BRCA (4.5%) |
| THYM | 15 | 0.867 | LUSC (6.7%), BRCA (6.7%) |
| SKCM | 12 | 0.917 | BRCA (8.3%) |
| BRCA | 133 | 0.925 | THCA (2.3%), PRAD (1.5%), CESC (0.8%) |
| KIRC | 62 | 0.935 | LIHC (3.2%), BRCA (3.2%) |
| SARC | 32 | 0.938 | TGCT (3.1%), KIRC (3.1%) |
| OV | 52 | 0.942 | UCEC (3.8%), TGCT (1.9%) |

| | | | |
|---|---|---|---|
| COAD | 54 | 0.944 | UCEC (3.7%), BRCA (1.9%) |
| THCA | 63 | 0.952 | BRCA (3.2%), LUSC (1.6%) |
| PCPG | 23 | 0.957 | TGCT (4.3%) |
| CHOL | 4 | 1.000 | — |
| LGG | 64 | 1.000 | — |
| LAML | 22 | 1.000 | — |
| PRAD | 62 | 1.000 | — |
| TGCT | 17 | 1.000 | — |
| UVM | 10 | 1.000 | — |

**Supplementary Table 2. Per-cancer-type classification accuracy and dominant misclassification patterns on the TCGA test dataset.**
Each test sample was classified by assigning the majority cancer-type label of its GMM component (learned from the TCGA training data). Accuracy (Acc) is the fraction of test samples correctly classified within a given cancer type. "Predicted Cancer Type" lists the cancer type(s) most frequently predicted instead of the true label, with the cancer-type-specific proportion indicated. N Samples is the number of TCGA test samples of that cancer type.

**Supplementary Table 3**

| Model | 1 | 2 | 3a | 3b | 4a | 4b | 5 |
|---|---|---|---|---|---|---|---|
| **GMM** | | | | | | | |
| N comps (K) | 70 | 75 | 140 | 90 | 110 | 120 | 100 |
| GMM mean init | (4.0, 5.0) | (4.0, 5.0) | (5.0, 5.0) | (5.0, 5.0) | (5.0, 5.0) | (7.0, 5.0) | (5.0, 5.0) |
| GMM SD init | (0.03429, 1.0) | (0.03733, 1.0) | (0.02286, 1.0) | (0.02222, 1.0) | (0.031818, 1.0) | (0.029167, 1.0) | (0.016, 1.0) |
| Weight alpha | 1.5 | 1 | 1.5 | 1 | 1 | 1.5 | 1.5 |
| **Reps (L)** | 30 | 35 | 25 | 35 | 30 | 20 | 35 |
| **Decoder** | | | | | | | |
| Shared | 100, 100,100 | 100, 100, 100 | 100, 100, 100 | 100, 100, 100 | 100, 100, 100 | 100, 100 | 100, 100, 100 |
| mRNA | 100, 512 | 128, 256 | 100, 512 | 128, 512 | 100, 128 | 128, 256 | 100, 256 |
| miRNA | 100, 256 | 128, 256 | 100, 128 | 100, 256 | 256 | 128, 128 | 128, 128 |
| R init | 4 | 4 | 4 | 2 | 2 | 4 | 4 |
| PI init | 0.25 | 0.75 | 0.75 | 0.5 | 0.75 | 0.5 | 0.5 |
| Activation | leaky_relu | relu | leaky_relu | relu | relu | relu | relu |
| **Training** | | | | | | | |
| Optimizer | AdamW | AdamW | AdamW | AdamW | AdamW | AdamW | AdamW |
| Learning rate | dec: 1e-4, gmm: 1e-2 rep: 1e-2 | dec: 1e-4, gmm: 1e-2 rep: 1e-2 | dec: 1e-4, gmm: 1e-2 rep: 1e-2 | dec: 1e-4, gmm: 1e-2 rep: 1e-2 | dec: 1e-4, gmm: 1e-2 rep: 1e-2 | dec: 1e-4, gmm: 1e-2 rep: 1e-2 | dec: 1e-4, gmm: 1e-2 rep: 1e-2 |
| Weight decay | dec: 1e-4, gmm: 1e-4 rep: 1e-4 | dec: 1e-3, gmm: 1e-4 rep: 1e-3 | dec: 1e-4, gmm: 1e-4 rep: 1e-4 | dec: 1e-3, gmm: 1e-4 rep: 1e-3 | dec: 1e-4, gmm: 1e-4 rep: 1e-3 | dec: 1e-4, gmm: 1e-3 rep: 1e-4 | dec: 1e-4, gmm: 1e-4 rep: 1e-4 |
| Betas | (0.9, 0.999) | (0.9, 0.999) | (0.9, 0.999) | (0.9, 0.999) | (0.9, 0.999) | (0.9, 0.999) | (0.7, 0.9) |
| **N params** | 9,774,885 | 5,061,968 | 9,667,687 | 9,707,386 | 2,617,326 | 4,810,354 | 4,147,982 |

**Supplementary Table 3: Model architecture and training specification.**
For each miDGD model configuration, the table reports Gaussian mixture model (GMM) hyperparameters (number of components, initialization of means and standard deviations,

mixture weight prior), latent dimensionality (number of representations), decoder architecture (shared, mRNA and miRNA branches), dispersion and zero-inflation parameters initialization, activation functions, and training settings (optimizer, learning rates, weight decay, betas). The total number of trainable parameters is given for each model (excluding sample-specific latent representations). These parameters were found using Weight and Biases sweeps.
